# Emergence of selectivity and specificity in a coarse-grained model of the nuclear pore complex with sequence-agnostic FG-Nups

**DOI:** 10.1101/2023.07.27.550776

**Authors:** Manoj Kumar Patel, Buddhapriya Chakrabarti, Ajay Singh Panwar

## Abstract

The role of hydrophobicity of phenylalanine-glycine nucleoporins (FG-Nups) in determining transport of receptor-bound cargo across the nuclear pore complex (NPC) is investigated using Langevin dynamics simulations. A coarse-grained, minimal model of the NPC, comprising a cylindrical pore and hydrophobic-hydrophilic random copolymers for FG-Nups was employed. Karyopherin-bound receptor-cargo complexes (Kaps) were modeled as rigid, coarse-grained spheres without (inert) and with (patchy) FG-binding hydrophobic domains. With a sequence-agnostic description of FG-Nups and the absence of any anisotropies associated with either NPC or cargo, the model described tracer transport only as a function of FG-Nup hydrophobicity, *f*. The simulations showed the emergence of two important features of cargo transport, namely, NPC selectivity and specificity. NPC selectivity to patchy tracers emerged due to hydrophobic Kap-FG interactions and despite the sequence-agnostic description of FG-Nups. Further, NPC selectivity was observed only in a specific range of FG-hydrophobic fraction, 0.05 ≤ *f* ≤ 0.20, resulting in specificity of NPC transport with respect to *f*. Significantly, this range corresponded to the number fraction of FG-repeats observed in both *S. cerevisiae* and *H. sapiens* NPCs. This established the centrality of FG-hydrophobic fraction in determining NPC transport, and provided a biophysical basis for conservation of FG-Nup hydrophobic fraction across evolutionarily distant NPCs. Specificity in NPC transport emerged from the formation of a hydrogel-like network inside the pore with a characteristic mesh size dependent on *f*. This network rejected cargo for *f* > 0.2 based on size exclusion which resulted in an enhanced translocation probability for 0.05 ≤ *f* ≤ 0.20. Thus, the NPC model provides a template for designing synthetic, biomimetic nanopores for macromolecular separations with high degrees of selectivity and specificity.

## INTRODUCTION

The nuclear pore complex (NPC) is a large biological nanopore that controls nucleocytoplasmic transport across the nuclear envelope (NE) membrane in a eukaryotic cell (1). It possesses an octagonal symmetry in the NE plane with both NPC structure and function being conserved across all eukaryotic species, from yeast to vertebrates. With an internal diameter estimated to vary between 30 – 50 nm, the NPC is composed of large proteins called nucleoporins (Nups) that are arranged in octagonal symmetry about its axis (2–5). Nearly thirty different Nups serve as building blocks of the NPC(6). A large fraction of the NPC mass (≈ 30%) is comprised of intrinsically disordered FG-Nups containing multiple phenylalanine (F) and glycine (G) repeats. Natively unfolded terminal domains of FG-Nups (∼200 – 800 residues long) line the central channel of the NPC resembling a “polymer brush” architecture on the inner periphery of NPC (7, 8).

The most striking attribute of the NPC, critical to its function as the “gatekeeper” of the nucleus, is the high degree of selectivity with which it allows the transport of myriad proteins and mRNA between the nucleus and cytoplasm (9). The NPC’s remarkable selectivity emerges from a complex interplay of specific interactions between FG-Nups of the NPC and FG-binding domains on karyopherin (Kap) receptors bound to cargo proteins (10). In addition, a RanGTP/GDP cycle provides directionality to nucleocytoplasmic exchange of cargo through the NPC (8, 9, 11). Whereas, smaller cargo (< 40 kDa or < 6 nm) may diffuse passively through the NPC, transport of larger cargo (> 60 kDa) requires binding of Kap (or other transport receptors) to the cargo (9).

Although, the central role of Kap/FG-Nup interactions in NPC transport is widely recognized, mechanisms of NPC transport remain elusive (8). An important reason is the lack of biophysical measurements that can resolve FG-Nup dynamics and Kap-cargo complex transport at molecular length scales (9, 11). In this context, several transport models have attempted to provide a physical basis for NPC selectivity based on its structure and FG-Nup/cargo interactions. These models include, the selective phase model (12), virtual gate model (13), reduction-of-dimensionality (ROD) model (14), and the forest model (8). However, none of the models provide a comprehensive description of selective transport through NPC (15).

Such limitations set the stage for use of molecular simulations to describe NPC dynamics from a molecular perspective. All-atom molecular dynamics (MD) simulations have shown that arrays of Nsp1 FG-Nups form extended brush-like structures and associate with NTF2 transport receptors via distinct binding spots (16, 17). However, millisecond (ms) time-scales typically associated with NPC transport are not practical for fully-atomistic MD simulations (18). Coarse-grained (CG) simulations have highlighted the complex interplay between NPC structure, receptor-FG-Nup interactions and intricate cargo motions through the NPC (19–21). Simulation studies have emphasized the role of receptor-cargo and FG-Nups binding interactions in overcoming entropic barriers to NPC entry and subsequently facilitating cargo transport through the NPC. In addition, CG simulations have provided insights into the heterogenous organization of FG-Nups and the NPC energy landscape resulting from FG-Nup interactions (22, 23).

Most CG studies have retained individual residue identities in FG-Nup descriptions, often replicating known FG-Nup organization in a particular NPC (either yeast or human) (22, 24, 25). Accordingly, the emergent NPC selectivity is a result of a heterogeneous set of FG-Nups lining the central channel. However, a sequence-agnostic approach to understanding NPC selectivity is desirable in conceptualizing (and designing) synthetic NPC mimic platforms for high-throughput macromolecular separations and bio-pumping applications (26, 27). Previously, Fragasso *et al.* (25) demonstrated NPC selectivity by modelling all FG-Nups with averaged distributions of GLFG repeats. This is significant finding but the model description alludes to specific chemical identities for residues. Hence, there is need to understand the NPC selectivity using a reduced and simpler physical model that can describe relevant FG-Nup dynamics and cargo transport.

The aim of this study was to capture the central role of FG-Nup hydrophobicity alone in nucleocytoplasmic transport by constructing a minimal computational model of NPC inspired nanopore channel with varying hydrophobicity. Langevin dynamics simulations and a polymer-based model of FG-Nups was used to investigate the transport of spherical receptor-cargo complexes through a cylindrical pore of typical NPC dimensions. Using a random copolymer description for FG-Nups, we investigated the role of Nup hydrophobicity on internal NPC organization, NPC selectivity, and cargo transport through the NPC. Our model presented here is different from previous studies in a way that we treated FG-Nups as random copolymer chains rather than considering their full amino acid sequence heterogeneity (22, 28–31) or intermolecular cohesiveness (25). With this reductionist approach, we focused on studying the effect of FG-Nups’ conformational entropy and hydrophobic interactions in absence of other physical factors. Our research adds to our understanding of the physical basis for NPC selectivity, as well as holds a potential to provide a platform for custom design and shape modification of drug-delivery payloads intended for the nucleus, and eventually inspiring NPC-mimetic filtering systems for industrial use (26, 32–34).

## SIMULATION METHOD

A coarse-grained (CG) approach was used to describe the structure of the NPC, which was modelled as a cylindrical pore with polymer brushes grafted in the inner (Figure 1A). The cylindrical NPC pore was composed of CG “wall” beads. FG-Nups were modelled as linear, random copolymers and each FG-Nup molecule was represented by a freely-jointed bead-spring chain with *N* = 300 beads. Every chain was composed of both hydrophilic (type-1) and hydrophobic (F*x*FG) (type-2) beads which were randomly distributed along the chain contour (Figure S1). The number of hydrophobic beads on an FG-Nup was determined by the hydrophobic fraction, *f*. Finally, individual FG-Nups were end-grafted on the inner surface of the cylindrical pore in nine equally-spaced rings. Each ring of FG-Nups was composed of eight polymer chains, thus mimicking the NPC’s eight-fold symmetry. The details of the CG NPC model are shown in Figure S1.

**Figure 1:**
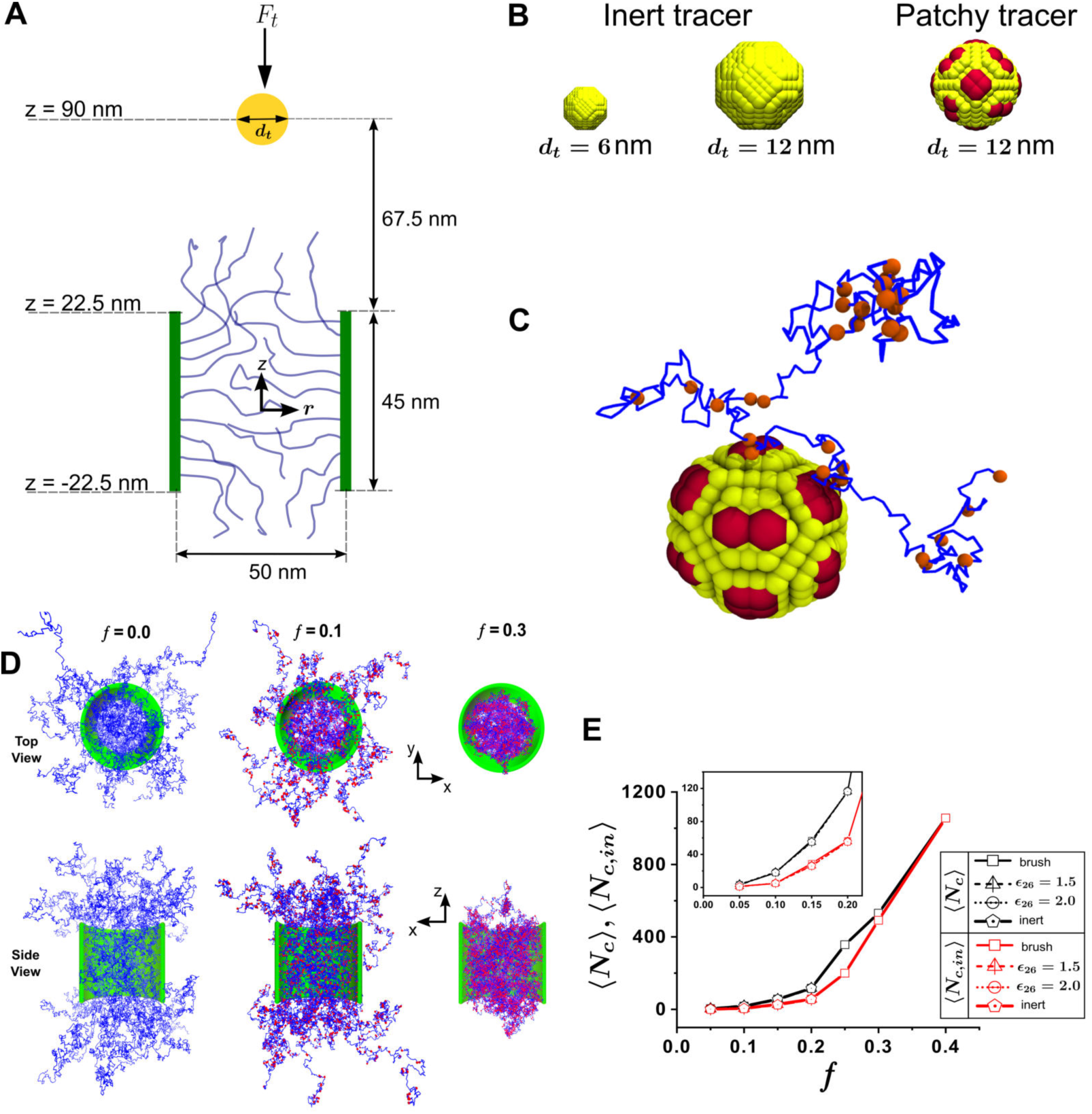
(A) Schematic of the simulation setup showing a spherical tracer placed outside a model cylindrical NPC pore with polymer brushes grafted on the inside periphery of the pore. A constant downward force of *F_t_* was applied on the tracer throughout the translocation process. (B) CG models of inert and patchy tracers of size, *d_t_*= 6 nm and 12 nm. Red colored patches on the patchy tracers represented the FG-binding pockets on the tracers. (C) A simulation snapshot of hydrophobic FG beads (also in a shade of red) on an FG-Nup (blue segments represented hydrophilic residues on the copolymer) binding to the FG-binding pocket of a patchy tracer. (D) Simulation snapshots showing top and side views of equilibrium brush structures for the model NPC corresponding to different values of *f* = 0, 0.1, 0.3. Brush collapse is evident from the NPC side view for *f* = 0.3. (E) Variation of the number of hydrophobic clusters in the entire brush, *N_c_* and inside the cylindrical region, *N_c_*_,*in*_, with *f*.

The FG-Nup bond length, *l*_0_, was used as the fundamental scale for length and was set to *l*_0_ = 1. Each 300 bead FG-Nup was modelled on Nsp1 (a 601 amino acid FG-Nup found in yeast), implying that every CG bead on a copolymer chain represented two contiguous residues (amino acids). Based on approximate residue sizes along a polypeptide chain, *l*_0_ ≈ 1 nm of length in real units. Bead diameters for both type 1 and type 2 beads on a copolymer chain (represented by blue and red colors, respectively, in Figure 1C,D) were chosen as *σ*_1_ = *σ*_2_ = 0.75*l*_0_, and the diameter of type 4 beads (pore wall) were set to *l*_0_. The diameter and length of the NPC cylindrical pore were chosen to be *d* = 50*l*_0_ and *L* = 45*l*_0_, respectively, which are in range of known NPC dimensions across different species.

Cargo particles, which represented translocating proteins attached to receptors, were modelled as hollow and rigid spherical particles composed of several CG beads. The constituent CG beads were of two types, type 5 (inert) beads shown as yellow beads in Figure 1B and type 6 (having affinity with type 2 or hydrophobic beads of FG-Nups) beads shown as red patches in yellow beads in Figure 1B. The affinity of type 6 beads with type 2 beads represented specific binding interactions of karyopherins with FG-repeats in NPC Nups (illustrated in Figure 1C). In the simulations, inert tracers were classified as cargo particles composed of only type 5 beads. In contrast, tracer particles composed of both types 5 and 6 beads were classified as “*patchy tracers*”. The surfaces of patchy tracers were marked by evenly spaced patches of type 6 beads which served as binding domains for FG-Nups during cargo translocation. Karyopherins, such as importin-β, are non-spherical and contain asymmetrically located FG-binding domains. Both factors are understood to play key roles in cargo reception and acceptance into the NPC (7, 28). The focus of the current study was to understand the role of FG-Nup hydrophobicity in tracer translocation. Hence, only spherical tracers with evenly spaced patches were considered to eliminate dependence on tracer shape and anisotropy. Both small and large tracers with diameters, *d_t_* = 6*l*_0_, 12*l*_0_, were considered in the simulations.

Implicit solvent Langevin dynamics was used to simulate tracer transport through the NPC. Details of the simulation method are provided in Section S1 of the SI. A constant, downward external force, 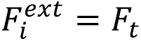, was applied to the tracer particle to nudge the tracer toward the NPC. This modeled the effect of Ran-GTP/GDP gradient in the biological scenario which confers directionality to cargo motion (4, 10, 24, 35–37). Although, application of a constant force appears to be an over-simplified representation of a more complex process, it is meant to mimic the essential features of directional nucleocytoplasmic transport in our simple CG polymer-based model.

Equations of motion were integrated in the NVE ensemble using the open-source MD package LAMMPS (38, 39). The Langevin piston was used to maintain the temperature at a constant value of 1 which can be mapped to a physiological temperature of 310 K. The NPC and the tracer were placed in a cuboidal simulation box of dimensions 100 × 100 × 250 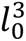 with fixed boundary conditions. The cylindrical pore extended from −22.5*l*_0_ ≤ *z* ≤ 22.5*l*_0_ in the *z*-direction. The dimensionless LJ interaction energies between pairs of hydrophobic-hydrophobic and hydrophilic-hydrophilic beads were chosen to be *ϵ*_22_ = 1.5 and *ϵ*_11_ = 0.1, respectively. These values have been shown to reproduce appropriate scaling behavior under varying solution conditions for a homopolymer (40)

The brush hydrophobic fraction was varied in the range *f* ∈ [0,0.4] to investigate the effect of FG content on tracer transport through the NPC. Prior to introducing the tracer, equilibrium configurations of FG-Nup brushes were generated for all hydrophobic fractions, *f* (Figure 1D and E). Equilibration times varied between 120,000 - 180,000 *τ*. After equilibration of the brush, tracer particles were introduced at *z* = 90*l*_0_, along the central axis of the pore, and were nudged toward the pore by applying a downward force, *F_t_*. Three values of *F_t_* = 0.2, 0.5, 1 (in reduced units) were considered, which are equivalent to 0.8 pN, 2 pN, and 4 pN, respectively. It is interesting to note that our choice of scales led to forces in the pN range which are the correct order of magnitude for forces in the biophysical context. To reiterate, the real world values the various characteristic scales in the simulations translate to as follows; *l*_0_ = 1nm, *T* = 310 K, *τ* = 20 ns, and *F* = 4 pN (see section S2 for a more detailed description of dimensionalization of characteristic scales in the simulation).

## RESULTS AND DISCUSSIONS

### Polymer Brush Density Distribution

Equilibrium structures of the model NPC for hydrophobic fractions, *f* = 0, 0.1 and 0.3, are shown in Figure 1D. Whereas, the brush remained extended outside the pore for lower values of *f* (< 0.25), it collapsed completely inside the pore for larger values of *f* (≥ 0.25). This behavior is evident from the top and side views shown in Figure 1D. Snapshots of equilibrium brush configurations for 0 ≤ *f* ≤ 0.4 (shown in Figure S2 of the Supporting Information) confirm the same behavior. An increase in *f* led to larger number of interactions between hydrophobic segments in the brush, resulting in formation of clusters of hydrophobic segments. Cluster formation in the brush is shown for *f* = 0.2 in Figure S3 of the Supporting Information. Hydrophobic clusters that formed due to attraction between hydrophobic segments were identified and quantified with the Stillinger algorithm (described in Section S2 of the Supporting Information) (41, 42). Two measures of cluster formation were calculated; (i) *N_c_*, the total number of hydrophobic clusters formed in the entire brush, and (ii) *N_c_*_,*in*_, the number of hydrophobic clusters that formed only within the cylindrical volume described by the pore. Both *N_c_* and *N_c_*_,*in*_ increased with *f* (Figure 1E) because of the increasing number of hydrophobic contacts between segments. Interestingly, *N_c_* was always larger than *N_c_*_,*in*_ for *f* < 0.3 because equilibrium brush configurations extended out of the cylindrical pore for lower FG fractions. However, *N_c_*_,*in*_ ≈ *N_c_* for *f* ≥ 0.3, consistent with a brush structure that had completely collapsed inside the pore.

Time-averaged segment densities along the axial and radial directions were also calculated for different values of *f* (Figure 2 and Figure S4 of the Supporting Information). The axial density plots in Figure 2A define brush height (outside the pore) as a function of *f* (the cylindrical pore spans −22.5 < *z* < 22.5), and clearly show brush collapse for *f* ≥ 0.3. Radial densities inside the pore (Figure 2B) increased with *f* and showed a peak toward the pore walls. However, radial density distributions appeared more uniform at lower *f* (< 0.2) indicating a more uniform distribution of hydrophobic clusters in the cylindrical volume of the pore. Although, most of the clusters were confined toward the pore walls at high hydrophobic fractions, it did not indicate a depletion region closer to the axis of the pore.

**Figure 2:**
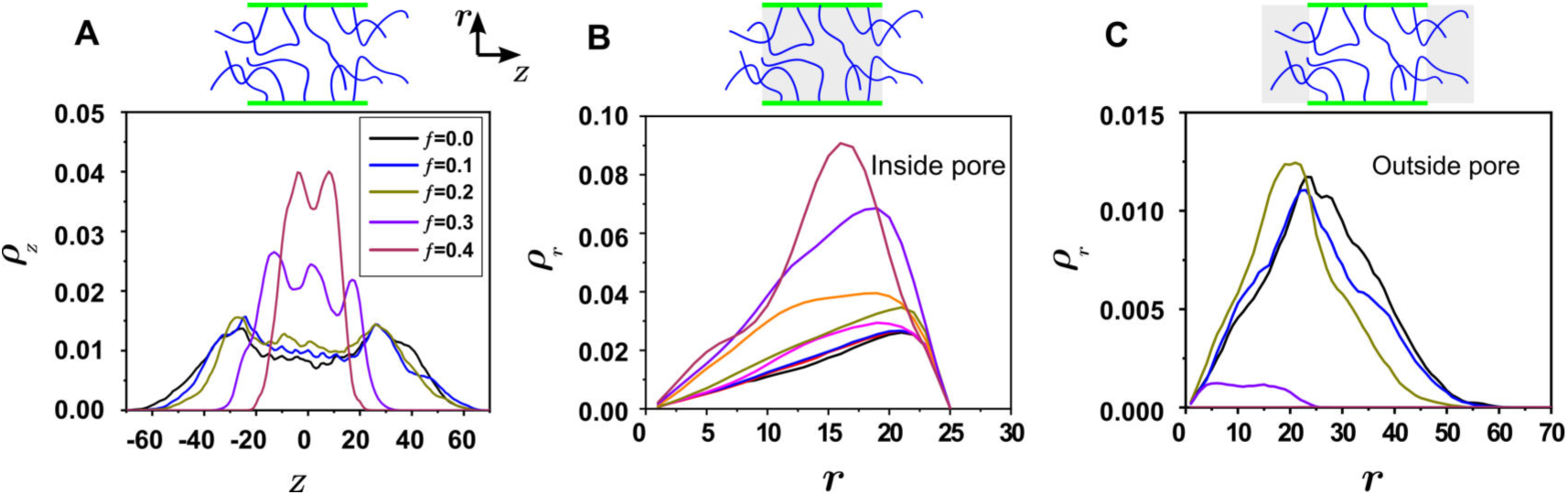
Plots showing equilibrated density distributions of brush segments along the (A) axial direction of the NPC, (B) radial direction of the NPC (inside the cylindrical region), and (C) radial direction of the NPC (outside the cylindrical region).

This was because segment densities were still higher compared to segment density values at lower hydrophobic fractions. A stark distinction between low and high hydrophobic fractions emerged in the radial density distributions calculated outside the pore (Figure 2C). The brush explored a large volume, described by a radius nearly twice that of the pore radius, for *f* ≤ 0.2. In contrast, the brush radial expanse outside the pore was practically non-existent for *f* > 0.2.

The change in equilibrium structure of the brush as a function of hydrophobic fraction is very important to the mechanisms of cargo transport across the model NPC. The extended brush at low *f* presents an entropic barrier to cargo entry and corresponds to the entropic repulsion model of NPC transport (13). In addition, an increase in *N_c_* (or *N_c_*_,*in*_) is indicative of a *gel-like network* formation in pore (see Figure S3 of the Supporting Information) and suggests that a selective-phase mechanism may be relevant with increasing hydrophobicity (11). Thus, our minimal coarse-grained model of the NPC demonstrates a malleable pore structure that can be altered from an entropic brush to a hybrid gel-brush to a tight network with the change of a single variable, *f*. This is of important biophysical consequence where the exact structure of the NPC is a topic of debate (43, 44).

### Tracer Trajectories

Cargo transport through the NPC was simulated by moving spherical tracer particles through the equilibrated brushes corresponding to different FG fractions. Cargo particles of two sizes, *d_t_* = 6 nm and 12 nm were chosen. Further, two types of tracers were simulated; (i) an *inert tracer*, which did not interact with the brush, and (ii) a *patchy tracer*, which had hydrophobic patches that could bind with the FG groups on the brush. Three different values of the downward force, *F_t_* = 0.8, 2 and 4 pN, respectively, were used in the simulations. Whereas, a total of ten independent trajectories were run for 6 nm tracers, twenty or more independent trajectories were simulated for 12 nm tracers. The simulated trajectories were classified into three types, *successful* (blue), *trapped* (pink) and *rejected* (black).

Inert tracers of *d_t_* = 6nm, corresponding to passive diffusion through the NPC, showed qualitatively different behaviors dependent on the value of *F_t_* (described in detail in SI). Tracer trajectories varied from absolutely no translocations at 0.8 pN, to few translocations at 2 pN and complete passage at 4 pN. Tracer trajectories for the larger, inert tracer (*d_t_* = 12 nm) at *F_t_*= 2 pN showed the first indications of NPC selectivity based on size exclusion (Figure S8 of the Supporting Information). Most of the trajectories were rejected for *f* = 0 and 0.05, consistent with the picture of entropic repulsion offered by the FG brush. This observation is in good agreement with several studies where inert cargoes are strongly barred to enter FG-permeable membranes hence their transport rate is extremely low (20, 25, 29, 31). Similarly, most trajectories were also rejected at higher hydrophobic fractions, *f* ≥ 0.25, because of the tight hydrophobic network which did not allow entry into the pore. Increased tracer translocation was detected for intermediate fractions, suggesting NPC selectivity for 0.05 < *f* < 0.25. One should be cautious in interpreting these results as evidence of NPC selectivity because these tracers were completely inert. It is well known that tracer transport through NPC is always facilitated by transport receptors. Hence, tracer transport should have been completely blocked by the NPC for an inert tracer, either by entropic repulsion at low *f* or by a tight mesh at high *f*. The observation of successful translocation was an artefact of our coarse-grained model where inert tracer particles interacted with the brush beads through purely repulsive Lennard-Jones interactions. With an additional downward pointing force, *F_t_*, this resulted in parting of the brush network and successful translocation attempts for a small number of trajectories in the inert tracer simulations.

The affinity of hydrophobic patches on patchy tracers was defined by the LJ interaction energy *ϵ*_26_ between FG groups (type 2 beads) and the patches on the tracers (type 6 beads). In the first case of patchy tracer simulations, *ϵ*_26_ = *ϵ*_22_ = 1.5, a notable increase in the number of successful translocations was observed when compared to the inert tracer (Figure 3B). Whereas, no translocations were observed for *f* ≥ 0.25, the pore was highly restrictive to tracer entry for lower fractions as well, *f* = 0 and 0.05. The maximum number of translocations occurred for *f* = 0.1. Thus, our coarse-grained model showed an enhanced selectivity of the NPC when the tracer contained hydrophobic patches that mimicked FG-binding domains on a cargo particle.

**Figure 3:**
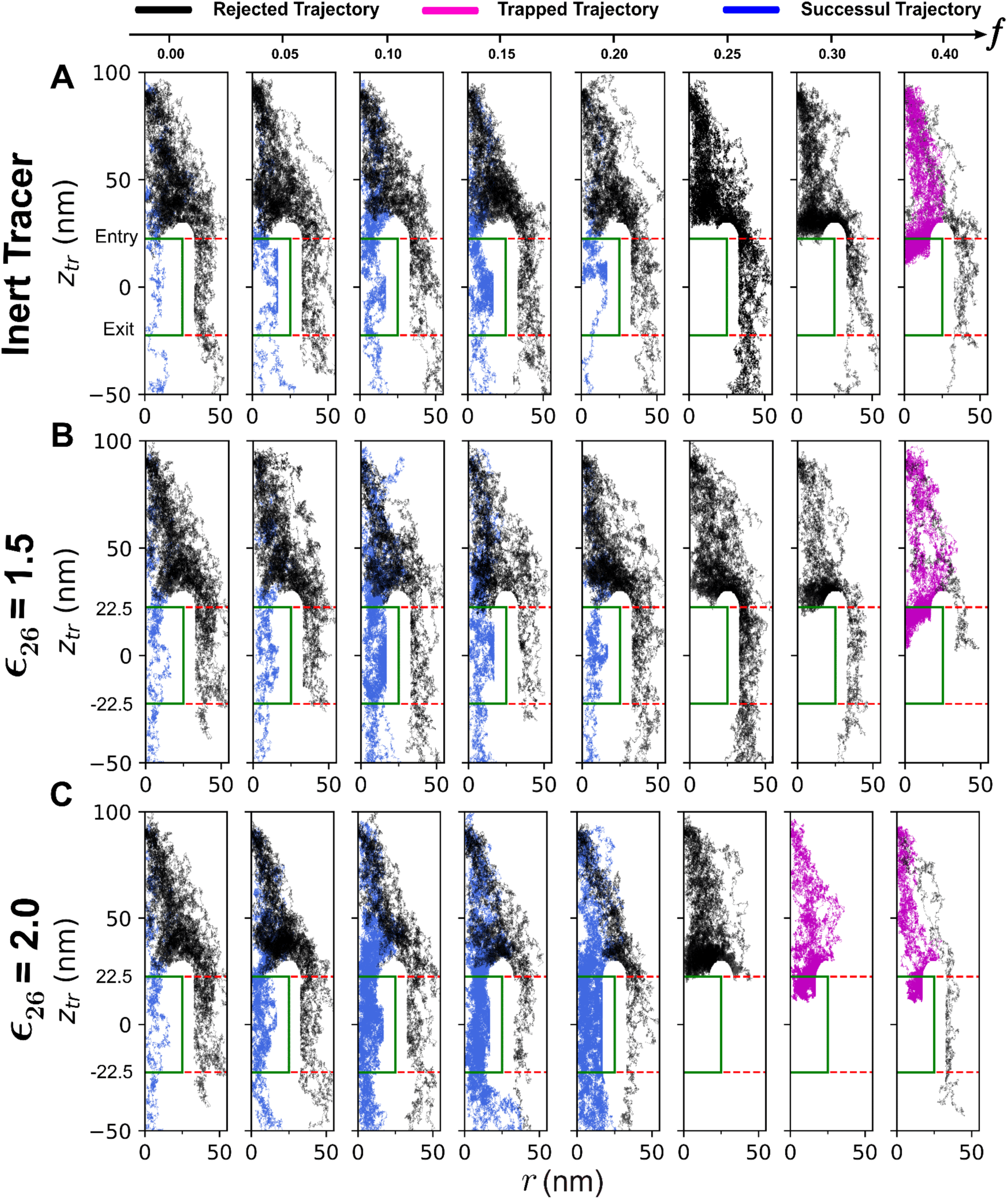
Tracer trajectories from twenty independent simulations shown for different values of *f* corresponding to (A) inert tracer of *d_t_*= 12 nm, (B) patchy tracer of *d_t_*= 12 nm, *ϵ*_26_ = 1.5, and (C) patchy tracer of *d_t_*= 12 nm, *ϵ*_26_ = 2.0, respectively. The trajectories show the variation of the *z*-coordinate of the tracer, *z_tr_*, with respect to the tracer radial coordinate, *r*. The value of *F_t_* was 2 pN for all three simulation sets. The simulated trajectories were classified into three types, *successful* (blue), *trapped* (pink) and *rejected* (black). The cylindrical pore is shown by a green rectangle from −22.5 ≤ *z* ≤ 22.5. Successful trajectories corresponded to tracers “successfully” entering the NPC and exiting from the other side.

It is well known that the Kap surface possesses greater hydrophobicity than other cytoplasmic proteins (45) and this hydrophobicity enables cargo to lower the entry barrier into the pore (46). Accordingly, a slight increase in the affinity of patches with hydrophobic beads in brush to *ϵ*_26_ = 2.0, led to a large increase in the number of successful translocations through the pore in the range 0.5 ≤ *f* ≤ 0.2 (see Figure 3). The most important observation from the tracer trajectories in Figure 3 was that tracer translocation through the NPC changed dramatically with both FG fraction, *f*, and the intensity of tracer-brush affinity (*ϵ*_26_). For *f* = 0, the entropic repulsion from the brush ensured near complete rejection of tracers from entry into the NPC. As stated above, the small fraction of trajectories (< 10%) that entered the NPC for *f* = 0, were an artefact of the simulation model. Interestingly, a previous study where phenylalanine residues (from Nsp1) were replaced by serine residues (Nsp1-S), also observed translocation of both inert cargo (tCherry) and patchy cargo (Kap95) pass through the NPC (29, 31). For higher FG fractions, *f* ≥ 0.25, the tight network of the NPC completely blocked tracer entry into the NPC. Thus, the NPC allowed tracer translocations only for intermediate FG fractions corresponding to 0.05 ≤ *f* ≤ 0.20, indicating that tracer-brush affinity is essential for NPC selectivity.

### Role of hydrophobicity

The importance of hydrophobic interactions between FG-binding domains on transport receptors and FG-Nups in enabling cargo transport through the NPC is well established. However, the importance of the number of FG-repeats in FG-Nups (*f*) and the intensity of FG-receptor hydrophobic interactions (*ϵ*_26_) are not well understood (7). It is believed that spatial segregation emerging from different FG-motifs is crucial in making the NPC selective for incoming cargo (22, 23, 25). The current study is unique because it attempts to demonstrate NPC selectivity emerging from a coarse-grained and minimal NPC model where hydrophobic FG repeats are placed randomly along FG-Nups. In other words, the model decouples sequence specificity of FG-Nups from cargo recognition and translocation through the NPC. Hydrophobic interactions between the FG segments resulted in the collective organization of Nups into qualitatively different equilibrium structures at different FG fractions, *f* (Figure 1D, Figure S2). In turn, these equilibrium structures resulted in selective tracer translocation through the NPC for a specific range of *f*. This leads to an important question; *What is the biophysical relevance of this narrow range of FG-hydrophobic fractions where the model NPC is selective to tracer transport?*

NPC functionality is known to be conserved across all eukaryotic species, including *Homo sapiens* and *Saccharomyces cerevisiae* (4, 36), even though NPC constituent elements may differ a lot for different species (47). The amino acid sequences for every FG-Nup for both *H. sapiens* and *S. cerevisiae* were studied by Peleg and Lim (48). The total number of FG-repeats present in every FG-Nup were extracted from their data. Then, the number fractions of FG-repeats in every FG-Nup were calculated and compared with the total number of residues in FG-Nups. Figure 4A shows these data for both *S. cerevisiae* and *H. sapiens*, respectively. These data were then plotted as histograms showing number fractions of FG-Nups with a specific percentage of FG-repeats for both species (Figure 4B). Interestingly, there was a close overlap of the two histograms which showed that most FG-Nups had FG-repeats in the range of 0.05 ≤ *f* ≤ 0.20. Only one human FG-Nup (Nup 214) had an FG-repeat fraction, *f* = 0.3, outside this range. This is an important observation because it suggests an important role for the number fraction of FG-repeats in NPC selectivity and transport.

**Figure 4:**
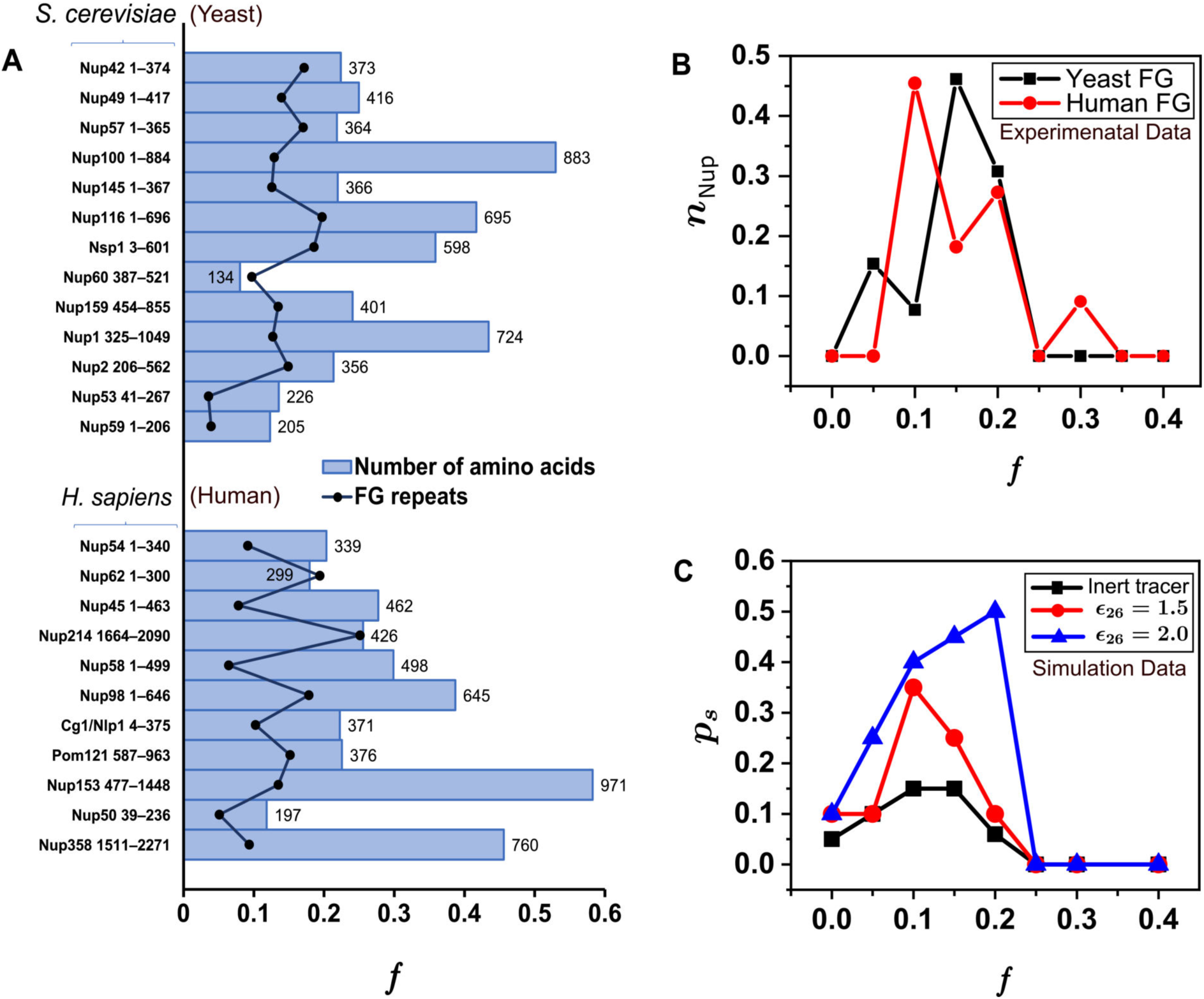
(A) Plots showing FG fraction and total number of residues of every FG-Nup for *S. cerevisiae* (yeast) and *H. sapiens* (human), respectively. These plots were generated based on data from Peleg and Lim (48). (B) Histogram of number fraction of FG-Nups, *n_Nup_*, plotted as a function of FG fraction, *f*, for both *S. cerevisiae* and *H. sapiens*, respectively. Maxima in *n_Nup_* were observed for 0.1 ≤ *f* ≤ 0.2 for both yeast and human Nups. (C) Probability of successful translocation, *p_s_*, plotted as a function of *f* for inert and patchy tracers corresponding to simulations with *d_t_*= 12 nm and *F_t_* = 2 pN. Maxima in *p_s_* were also observed for 0.1 ≤ *f* ≤ 0.2.

Further, the range of FG-repeats also coincides with the range of hydrophobic fractions that allowed most trajectories to pass through our modelled NPC pore in Figure 3. The probability of successful translocation of a tracer particle, *p_s_*, for a particular *f* was defined as,

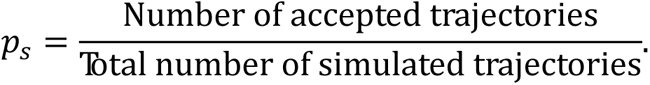

Figure 4C plots *p_s_* as a function of *f* for an inert tracer and patchy tracers (with *ϵ*_26_ = 1.5 and 2.0), respectively. A comparison of Figures 4B and 4C led to three important results related to NPC translocation. First, *p_s_* increased with increasing tracer-FG affinity (*ϵ*_26_). Second, the *p_s_* vs *f* plots clearly showed that the NPC was highly selective to tracer entry and translocation in a narrow range of FG-fractions, 0.05 ≤ *f* ≤ 0.20. Third, and most importantly, the range of FG-fractions where translocation was allowed corresponded to the population of FG-repeats observed in both *S. cerevisiae* and *H. sapiens* in Figure 4B. Thus, our reduced NPC model demonstrated both selectivity (recognition of transport-receptor bound cargo) and specificity (a narrow permissible range of FG-fractions) that are hallmarks of nucleocytoplasmic transport through the NPC.

The *p_s_* values for inert tracers remained very low, with a maximum of ≈ 0.15 at *f* = 0.10. In comparison, for patchy tracers, more than two-fold (for *ϵ*_26_ = 1.5) and more than three-fold (for *ϵ*_26_ = 2.0) increase in the maximum *p_s_* values were observed, respectively. The choice of a higher tracer-brush interaction strength was inspired by previous observations that FG-FG hydrophobic interaction is generally weaker in comparison to the collective hydrophobicity of Kaps, so that FG-Nups can differentiate between FG-FG and FG-Kaps interaction (20, 28, 45, 49, 50). When compared with translocation at *ϵ*_26_ = 1.5, *p_s_* values increased over the permissible range for *ϵ*_26_ = 2.0 and the maximum shifted to *f* = 0.20. This can be attributed to enhanced tracer-FG-Nup hydrophobic interactions at *ϵ*_26_ = 2.0, which led to a dramatic five-fold increase in *p_s_* at *f* = 0.20 (from *p_s_* = 0.1 at *ϵ*_26_ = 1.5 to *p_s_* = 0.5 at *ϵ*_26_ = 2.0). According to the selective phase model, Kap can locally break down network crosslinks during cargo translocation because Kap-FG affinity is higher than FG-FG affinity (7, 22). The dynamic modification of the FG-Nup network helps the Kap-cargo complex move through the sieve-like gel. The large increase in *p_s_* values at *ϵ*_26_ = 2.0 appeared to be consistent with this view and reinforced the importance of FG-binding domains in Kaps in determining NPC selectivity.

All translocations were completely blocked for *f* ≥ 0.25 for all types of tracers, highlighting the selective nature of the model NPC. At *f* = 0, *p_s_* values were very small (≤ 0.1) for all types of tracers, indicating that the expanded FG-Nup brush repelled tracers successfully in the absence of any tracer-FG-Nup interactions. An intermediate FG-fraction range, 0 < *f* < 0.25, represents the possibility of increased tracer-FG-Nup interactions which could result in enhanced *p_s_* for this range of *f*. The non-monotonic *p_s_* vs *f* curves in Figure 4C conclusively demonstrate that NPC selectivity increases with increasing tracer-FG-Nup interactions (*f*) before decreasing dramatically to zero for higher FG-Nup fractions (*f* ≥ 0.25). In the biophysical context, this narrow permissible range of FG-Nup fractions represents the specificity of the NPC when compared with the population of FG-repeats in both yeast and humans.

In a previous work, Fragasso *et al.* (25, 30) reconstituted NPC selectivity using GLFG-inspired artificial Nups, but did not relate NPC selectivity to hydrophobic content of the brush. In another simulation study, Ghavami *et al.* showed that effective cargo translocation is enabled by both the overall number of FG-binding domains on Kaps (number of patches) and their density distribution on the transport receptor (28). The same study and other simulations (28, 51) have also shown that a highly localized distribution of FG-binding sites on the transport receptor resulted in more selective translocation. This was found to be consistent with the highly-localized FG-binding domains found on Kaps (7). Our study is distinct from these previous investigations in two important ways. First, by considering a spherical tracer with an isotropic distribution of hydrophobic patches (FG-binding domains), it decouples FG-binding domain distribution on the tracer from NPC selectivity. Second, the results from Figure 4C demonstrate that NPC selectivity can be mimicked in a minimal, coarse-grained model by simply changing the hydrophobic fraction, *f*, in a random copolymer (“FG-Nup”). In contrast with studies which considered specific FG-Nup sequences (22–25, 31), the current model mimics both NPC selectivity and specificity in a sequence-agnostic manner by considering hydrophobic beads of equal hydrophobicity in an FG-Nup.

### Hydrophobic Clusters and Mesh Formation

Hydrophobic clusters, that formed due to association of hydrophobic FG beads on the FG-Nups (discussed in Figure 1E), were used to quantitatively describe network formation inside the pore as a function of *f*. Given the highly dynamic nature of the brush, both composition of individual clusters (in terms of the constituent beads) and the number of clusters, *N_c_*, were dynamic with respect to time. However, equilibrium values of both 〈*N_c_*〉 and 〈*N_c_*_,*in*_〉 depended only on *f*. Two length scales were defined to describe the network. The equilibrium mesh size of the pore, 〈*ξ*〉, was defined as the average distance between four closest hydrophobic clusters (network points) that formed a “pore” in the network (Figure 5A). The final value calculated at a particular FG fraction, *f*, represented an average computed over the entire pore and over multiple time frames of the trajectory. A second length scale, the peripheral cluster distance, 〈*r_p_*〉, was also calculated in a similar way. This represented the average size of the opening in the network between peripheral clusters and the pore walls (Figure 5B). Mechanistically, 〈*ξ*〉 and 〈*r_p_*〉, represented two different paths that a tracer could follow while translocating through the NPC. Whereas, 〈*ξ*〉 corresponded to the tracer moving through the network closer to the pore axis, 〈*r_p_*〉 corresponded to a tracer moving along the openings in the network closer to pore walls. Both pathways have been proposed previously in models of NPC transport.

**Figure 5:**
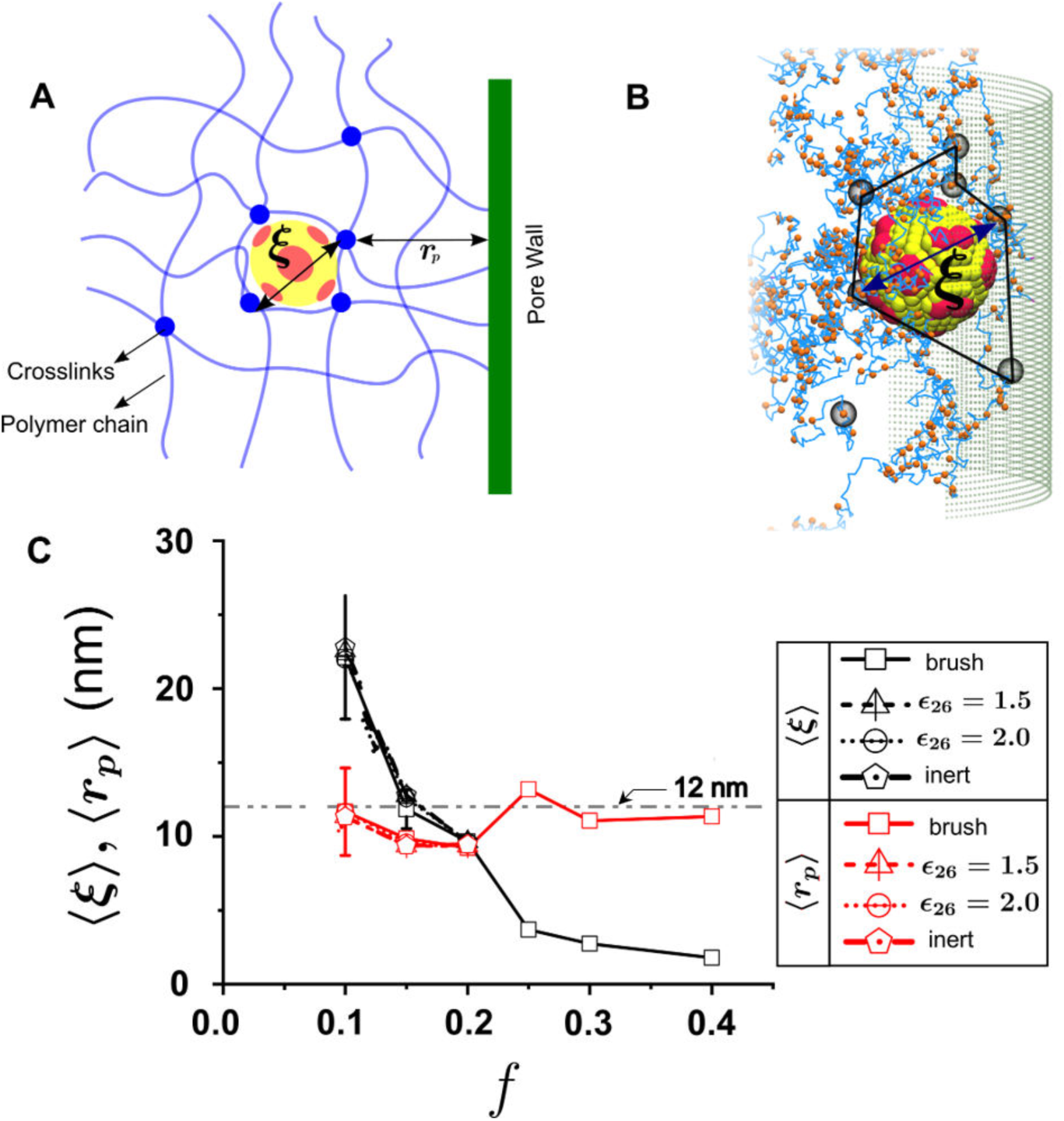
Mesh and cluster formation inside pore. (A) A schematic illustration of a patchy tracer interacting with FG-Nups that form a cross-linked mesh network inside the NPC. Two characteristic length scales of the network, the mesh size, *ξ*, and the peripheral distance of crosslinks (or clusters) from pore wall, *r_p_*, are shown also shown. (B) A simulation snapshot showing the formation of a mesh defined by network points of hydrophobic clusters (originating from FG-FG interactions) and a patchy tracer (*d_t_*= 12 nm, *ϵ*_26_ = 1.5, *f* = 0.15) residing within this. (C) The average values of mesh size, 〈*ξ*〉, and peripheral distance, 〈*r_p_*〉, plotted as functions of *f*. Both length scales were calculated for the cases of only the equilibrated brush (no tracer present) and during tracer translocation.

The changes in 〈*ξ*〉 and 〈*r_p_*〉 with respect to *f* are plotted in Figure 5C in absence and presence of a tracer. Interestingly, 〈*ξ*〉 and 〈*r_p_*〉 values in the pore remained unaffected by the presence of a tracer, and were found to be unchanged even when different tracer types were considered (implying varying levels of brush-tracer interactions, *ϵ*_26_). This indicated that the tracer did not disturb the averaged network-structure significantly during translocation through the pore.

Figure 5C shows that significant network formation inside the pore occurred only for *f* ≥ 0.1, with 〈*ξ*〉 ≈ 22 nm at *f* = 0.1. A rapid decrease in 〈*ξ*〉 was observed with increase in *f* with 〈*ξ*〉 ≈ 2 nm for *f* = 0.4. In contrast, 〈*r_p_*〉 values remained nearly constant with *f*, varying in a narrow band between 10 – 12 nm. A dashed horizontal drawn at 12 nm in Figure 5C indicates the tracer size, and clearly shows that 〈*ξ*〉 is significantly smaller than the tracer size for *f* > 0.2. This correlates with the precipitous drop in *p_s_* values for *f* > 0.2 (Figure 4C) and leads to a size exclusion effect due to a much tighter network for *f* > 0.2. Thus, the network description of the pore clearly shows that tracer translocation is possible only in a narrow band, 0.5 ≤ *f* ≤ 0.2, where both 〈*ξ*〉 and 〈*r_p_*〉 are large enough to allow tracers to pass through the network.

### Time scales of tracer translocation

The time for tracer translocation was characterized by defining three different time-scales, namely, (i) an approach time, *τ_a_*, (ii) an interaction time, *τ_i_*, and (iii) a transit time, *τ_t_* (shown schematically in Figure 6A). All tracers started at *z* = 90 and *τ_a_* was calculated as the time taken by the tracer to move from *z* = 90 to *z* = 〈ℎ〉, where 〈ℎ〉 is the brush height at a specific FG-fraction *f*. The time interval from the tracer’s first contact with the brush at *z* = 〈ℎ〉 and its entry into the pore at *z* = 16.5 was defined as *τ_i_*. Finally, the time interval from tracer entry at *z* = 16.5 and its complete exit corresponding to *z* = −28.5 was defined as *τ_t_*.

**Figure 6:**
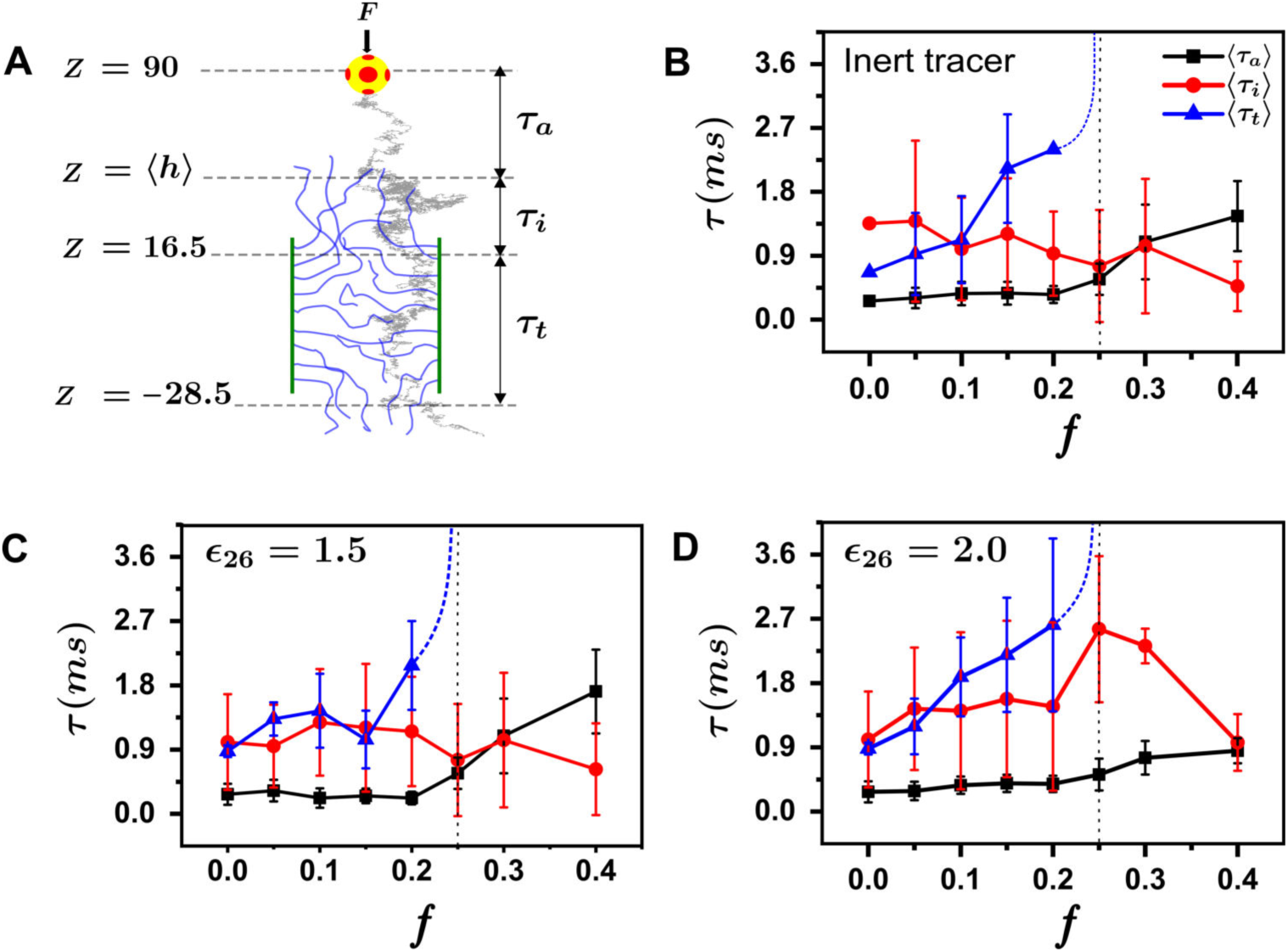
(A) Schematic defining the three calculated time scales in the simulations, namely, approach time (*τ_a_*), interaction time (*τ_i_*) and transit time (*τ_t_*). A typical tracer trajectory that successfully translocates through the pore is shown in grey. The variation of *τ_a_*, *τ_i_* and *τ_t_* with respect to brush hydrophobic fraction, *f*, for (B) an inert tracer, (C) a patchy tracer with *ϵ*_26_ = 1.5, and (D) a patchy tracer with *ϵ*_26_ = 2.0.

The changes in all three time-scales with respect to *f* for the three tracer types, inert, patchy (*ϵ*_26_ = 1.5) and patchy (*ϵ*_26_ = 2.0) are shown in Figures 6 B – D, respectively. Very small variations in *τ_a_* were observed for *f* ≤ 0.2 for all three tracer types, and *τ_a_* values increased for *f* > 0.2, consistent with brush collapse for higher FG-fractions. As expected, *τ_i_* behavior depended on the strength of tracer-brush interactions. For inert tracers, *τ_i_* decreased slightly with *f* because brush height decreased with *f*, indicating that tracer-brush interaction depended only on the spatial extent of the brush. For patchy tracers, *τ_i_* increased sharply when *ϵ*_26_ increased from 1.5 to 2.0. The increase was most noticeable for higher FG-fractions (*f* > 0.1) with *τ_i_* showing a maximum at *f* = 0.25 for *ϵ*_26_ = 2.0. This happened because of stronger tracer-brush interactions at *ϵ*_26_ = 2.0 and resulted in increased tracer entry into the pore. Enhanced tracer-brush interactions further led to a significant increase in *p_s_* for *f* = 0.15, 0.2 at *ϵ*_26_ = 2.0 (Figure 4C). This was observed as the shift in the location of the *p_s_* peak in Figure 4C from *f* = 0.1 at *ϵ*_26_ = 2.0 to *f* = 0.2 at *ϵ*_26_ = 2.0. Analysis of radial distribution of tracer positions, *ρ_tr_*, outside the pore (before entry) only for those tracer trajectories that successfully passed through the pore reinforced the link between a large *τ_i_* (at higher *ϵ*_26_) and an increase in *p_s_*. Figure 7 shows a comparison of *ρ_tr_* vs. *r* plots for different tracer-brush interactions at different *f*. Shifts toward larger values of *r* were observed with increase in *ϵ*_26_ for all *f*, indicating that a greater number of off-axis tracer trajectories successfully entered the pore. In contrast, inert tracer trajectories needed to be closer to the pore axis for successful translocations. These shifts were most prominent at larger FG-fractions of *f* = 0.15, 0.2, indicating that the probability of tracer capture by the brush depended strongly on both *f* and *ϵ*_26_. Consequently, higher tracer capture probabilities resulted in higher probabilities of tracer entry into the pore and successful translocations.

**Figure 7:**
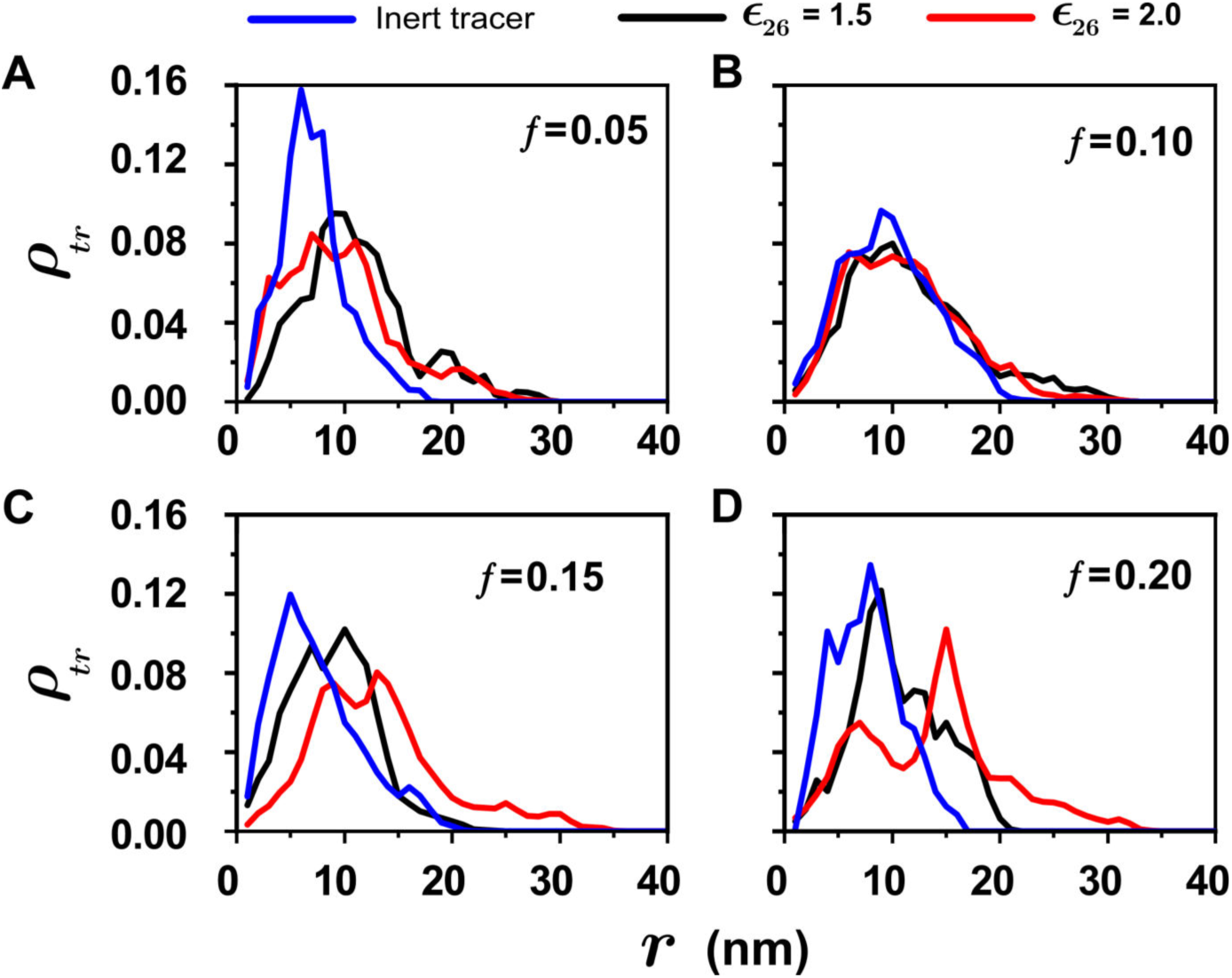
Radial density distribution of tracer positions, *ρ_tr_*, outside the pore (before entry) plotted only for successful tracer trajectories corresponding to different hydrophobic fractions, (A) *f* = 0.05, (B) *f* = 0.10, (C) *f* = 0.15, and (D) *f* = 0.20. Radial density distributions are shown for all three tracer types including, inert tracers and both types of patchy tracers with *ϵ*_26_ = 1.5 and 2.0.

Finally, *τ_t_* values increased monotonically over the range, 0 < *f* < 0.25, for all tracer types, asymptotically approaching *τ_t_* → ∞ at *f* = 0.25 (no tracer entry and translocation for *f* > 0.2). The monotonic increase in *τ_t_* with respect to *f* was due to two reasons; (i) decrease in mesh size, 〈*ξ*〉, with increasing *f*, and (ii) increase in tracer-network interactions with increasing *f*. A careful observation of the time axes labels in Figure 6 shows that all three time-scales are on the order of few milliseconds. Dimensional conversion of the inherent dimensionless simulation time-scales was carried out with appropriate physical scales (see Methods section). For instance, the total translocation time (*τ_a_* + *τ_i_* + *τ_t_*) for a patchy tracer at *f* = 0.2 and *ϵ*_26_ = 2.0, was approximately 5 ms. Experimentally determined NPC translocation time-scales for protein cargo are in a similar range, indicating that our coarse-grained and highly reduced NPC model can reproduce physically relevant time-scales (4, 52, 53). Significantly, it shows that the simplified copolymer description of the FG-Nups captures the most relevant physical aspects of protein translocation through the NPC.

## CONCLUSIONS

Coarse-grained Langevin dynamics simulations were used to examine the crucial role of FG-Nup hydrophobicity in cargo transport through a model NPC. A minimal model was adopted with; a cylindrical pore for the NPC, a random copolymer brush to represent FG-Nups and spherically-shaped receptor-cargo complexes. Incorporating a sequence-agnostic (random hydrophilic-hydrophobic copolymer) description of FG-Nups and in absence of any anisotropies associated with either NPC or cargo, the model described tracer transport only as a function of FG-Nup hydrophobicity.

Despite the highly reduced description, the model successfully captured the emergence of two important features of NPC transport, namely, (a) *selectivity*, involving selective passage of only receptor-bound cargo with FG-binding domains through the NPC, and (b) *specificity*, where cargo transport was allowed only over a narrow range of FG-Nup hydrophobicity. The model NPC was highly selective to patchy tracers (cargo with FG-binding domains) and showed a pronounced increase in *p_s_* (probability of successful translocation) values over inert tracers (Figures 3 and 4C). It was interesting that NPC selectivity emerged despite assuming no variations in either FG-Nup molecular weight or FG-Nup hydrophobic fraction, *f*, through the NPC. This novel result shows that the physical mechanism of NPC selectivity can be described in a sequence-agnostic manner, based only on overall FG-Nup brush hydrophobicity.

The model also showed that the transport of receptor-bound cargo across the NPC occurred only over a specific (and narrow) range of FG-fractions, 0.05 ≤ *f* ≤ 0.20. Remarkably, this mimicked, very closely, the number fraction of FG-repeats observed in both *S. cerevisiae* (yeast) and *H. sapiens* (human) (Figure 4). The most important finding of the current work is in providing a physical basis for the observed number fraction of FG-repeats in both yeast and human NPCs. Further, the result also provides a biophysical basis for conservation of FG-Nup hydrophobic fraction across evolutionarily distant NPCs. To the best of our knowledge, this is the first study of NPC transport that highlights the central role of FG-hydrophobic fraction in controlling NPC transport.

NPC specificity originated from the organization of FG-Nups inside the NPC. FG-FG interactions resulted in the formation of a hydrogel-like network with a characteristic mesh size that was dependent on the FG-hydrophobic fraction (Figure 5). For higher hydrophobic fractions (*f* > 0.20), the mesh size was significantly smaller than the cargo size. Consequently, passage of cargo through the NPC was determined by a size exclusion effect which led to enhanced translocation probability in a specific range of FG-fractions (0.05 ≤ *f* ≤ 0.20). To the best of our knowledge, the current study is the first which provides a quantitative description of a three-dimensional hydrogel-like network inside the NPC. In addition, FG-Nups formed an extended brush configuration outside the cylindrical pore for this range of *f*. Whereas, extended brush configurations presented an entropic barrier to entry of inert cargo, they assisted in the capture of receptor-bound cargo through attractive interactions between FG-binding domains and FG-Nups. Thus, a hybrid transport mechanism, combining both virtual gate and selective-phase (hydrogel) models, determined cargo transport through the NPC in the relevant (allowed) range FG-hydrophobic fraction (0.05 ≤ *f* ≤ 0.20). Tracer translocation time-scales were on the order of a few milliseconds (Figure 6) which agreed with typical time-scales observed in experiments. Therefore, the highly reduced NPC model was able to faithfully capture the most significant aspects of receptor-bound cargo transport through the NPC.

However, receptor-bound cargo transport through the NPC is a very complex process which is bound to be influenced by several factors, including residue-specific interactions, NPC structure, cargo shape and location of FG-binding domains on receptors. Some of these effects can be considered within the scope of the current model, and we are currently investigating effects of NPC shape anisotropy and FG-Nup molecular weight. We anticipate that these factors may modulate the current results related to *p_s_* and mesh organization. The coarse-grained model proposed here provides a template to reconstruct (or reproduce) NPC-like selectivity in a biomimetic in-vitro platform without adopting specific FG-Nup sequences. Finally, the minimal model is not limited to reproducing nucleocytoplasmic transport. It may guide design of devices, such as a highly-selective molecular filter (54, 55) with synthetic polymers playing the role of FG-Nups.

## SUPPORTING MATERIAL

Electronic Supporting Information, simulation movies and visualization.

## AUTHOR CONTRIBUTIONS

M.K.P. and A.S.P. designed the research and built systems. M.K.P performed simulations, analysed data, created figures, and prepared the initial manuscript draft. All authors contributed to the writing and editing of the final manuscript.

## ACKNOWLEDGEMENTS

We would like to acknowledge support in the form of access to various high-performance computing resources. These include the Dendrite high-performance computing facility in the Department of Metallurgical Engineering and Materials Science, IIT Bombay and Spacetime High Performance Computing (HPC) resource at IIT Bombay.

