## Supplementary Information for "Emergence of selectivity and specificity in a coarse-grained model of the nuclear pore complex with sequence-agnostic FG-Nups"

**Table of Contents** **Page**


| | | |
|---|---|---|
| **S1. Langevin dynamics simulations** | | S3 |
| **S2. Langevin dynamics simulation scales** | | S5 |
| **Figure S1:** | Schematic of Simulation Setup | S6 |
| **Figure S2:** | Snapshots of equilibrated brush for hydrophobic fractions, $f$=0 to 0.4 | S7 |
| **S3. Cluster formation algorithm** | | S8 |
| **Figure S3:** | Cluster identification algorithm | S8 |
| **Figure S4:** | Snapshot of clusters formation inside the pore for FG-fraction, $f$=0.2 | S9 |
| **Figure S5:** | Plots showing equilibrated density distributions of brush segments | S10 |
| **S4. Tracer trajectories of 6nm inert tracer** | | S11 |
| **Figure S6:** | Inert tracer trajectory at $F$=0.8pN for tracer size, $dt$ =6 nm w.r.t (A) radial direction, $r$ (B) time, $t$ | S12 |
| **Figure S7:** | Inert tracer trajectory at $F$=4pN for tracer size, $dt$ =6 nm w.r.t (A) radial direction, $r$ (B) time, $t$ | S13 |
| **Figure S8:** | Inert tracer trajectory at $F$=2pN for tracer size, $dt$ =6 nm w.r.t (A) radial direction, $r$ (B) time, $t$ | S14 |
| **S5. Tracer trajectories for inert and patchy tracers** | | S15 |
| **Figure S9:** | Inert tracer trajectory at $F$=2pN, $dt$ =12 nm w.r.t time, $t$. | S15 |
| **Figure S10:** | Patchy tracer trajectory at $F$=2pN, $dt$ =12 nm, $\epsilon_{26}$ = 1.5 w.r.t time, $t$. | S15 |
| **Figure S11:** | Patchy tracer trajectory at $F$=2pN, $dt$ =12 nm, $\epsilon_{26}$ = 2.0 w.r.t time, $t$. | S16 |
| **SI References** | | S17 |

### S1. Langevin dynamics simulations

All CG atoms in the simulation were assumed to execute Brownian motion in an implicit solvent. Their dynamics was described by the Langevin equation, S1 (1, 2) ,

$$m_i \frac{d^2 r_i}{dt^2} = -\zeta v_i - \nabla_i U(r_{ij}) + F_i^r(t) + F_i^{ext} \qquad \text{S1}$$

where, $m_i$, $r_i$, $\zeta$, and $v_i$ represented mass, position, damping coefficient and velocity of any particle $i$, respectively. The only exceptions were the CG atoms that made up the wall of the cylindrical pore, which were assumed to be immobile during the simulation. Random forces, $F_i^r$, acting on CG atoms were described by the fluctuation-dissipation theorem described in S2,

$$\langle F_i^r(t) F_j^r(t') \rangle = 6\zeta k_B T \delta(t - t') \delta_{ij}, \qquad \text{S2}$$

A constant, downward external force, $F_i^{ext} = F$, was applied to the tracer particle to nudge the tracer toward the NPC. This models the effect of Ran-GTP/GDP gradient in the biological scenario which confers directionality to cargo motion (3–8). Although, the application of a constant force appears to be an over-simplified representation of a more complex process, it is meant to mimic the essential features of directional nucleocytoplasmic transport in our simple CG polymer-based model.

The net interaction potential, $U(r_{ij})$ was described as a sum of excluded volume interactions, $U_{LJ}$, and bond potentials, $U_{bond}$ (for the copolymer chains) as shown in equation S3,

$$U(r_{ij}) = U_{LJ} + U_{bond}, \qquad \text{S3}$$

The excluded volume interactions between any two CG beads were described by the Lennard-Jones interactions given by equation S4,

$$U_{Lj}(r_{ij}) = \begin{cases} 4\epsilon_{LJ}\left[\left(\frac{\sigma}{r_{ij}}\right)^{12}\right] - \left[\left(\frac{\sigma}{r_{ij}}\right)^{6}\right], & r_{ij} \leq 2.5\sigma \\ 0, & r_{ij} > 2.5\sigma \end{cases} \qquad \text{S4}$$

where, $\epsilon_{LJ}$ and $\sigma$ are the Lennard-Jones interaction parameter and van der Waals diameter, respectively. Bond potentials, $U_{bond}$, for freely-jointed copolymer chains were described by the FENE (finitely-extensible non-linear elastic) potential (9, 10) as mentioned in equation S5,

$$U_{bond} = -\frac{1}{2}kR_{max}^2 \ln[1 - (r_b/R_{max})^2] \qquad \text{S5}$$

where, maximum allowable distance between beads $R_{max} = 1.5l_0$, $r_b$ is separation distance between any adjacent beads, $l_0$ is the bond length along the polymer chain and $k$ is spring constant. The velocity-Verlet algorithm (2) was employed to solve equation 1 numerically with a timestep $\Delta t$ of $0.001\tau$. Since, the Langevin equation was solved in the over-damped regime, the corresponding scale for time was described by $\tau = \zeta l_0^2/k_B T$s, where the Stokes drag coefficient is $\zeta = 6\pi\eta\sigma$. Correspondingly, the thermal energy, $k_B T$, was the scale for energy and $k_B T/l_0$ was the appropriate scale for force in the simulations.

### S2. Langevin dynamics simulation scales

The dimensionless simulation scales correspond to actual dimensional values in the following manner;

i. The scale for length, $l_0$, corresponds to approximately the size of two adjacent residues along a polypeptide chain, which is approximately 1 nm. Hence,

$$l_0 = 1 \equiv 1 \text{ nm}$$

ii. All simulations are relevant at the physiological temperature of 310 K. Therefore,

$$T = 1 \equiv 310 \text{ K}$$

iii. Thermal energy is used as the relevant scale for energy. Hence,

$$\epsilon = 1 \equiv k_B T$$

iv. Accordingly, force is defined as,

$$F = 1 \equiv \frac{\epsilon}{l_0} \equiv \frac{k_B T}{l_0} \equiv 4 \text{ pN}$$

v. A characteristic time-scale corresponding to a diffusion time is defined as,

$$\tau = 1 \equiv \frac{6\pi\eta a}{K_B T} l_0^2 = 22 \text{ ns}$$

which assumes a cellular viscosity of $\eta = 5$ cP (11, 12). However, this value was rounded off to $\tau = 20$ ns, which was chosen as the characteristic time-scale in the simulations.

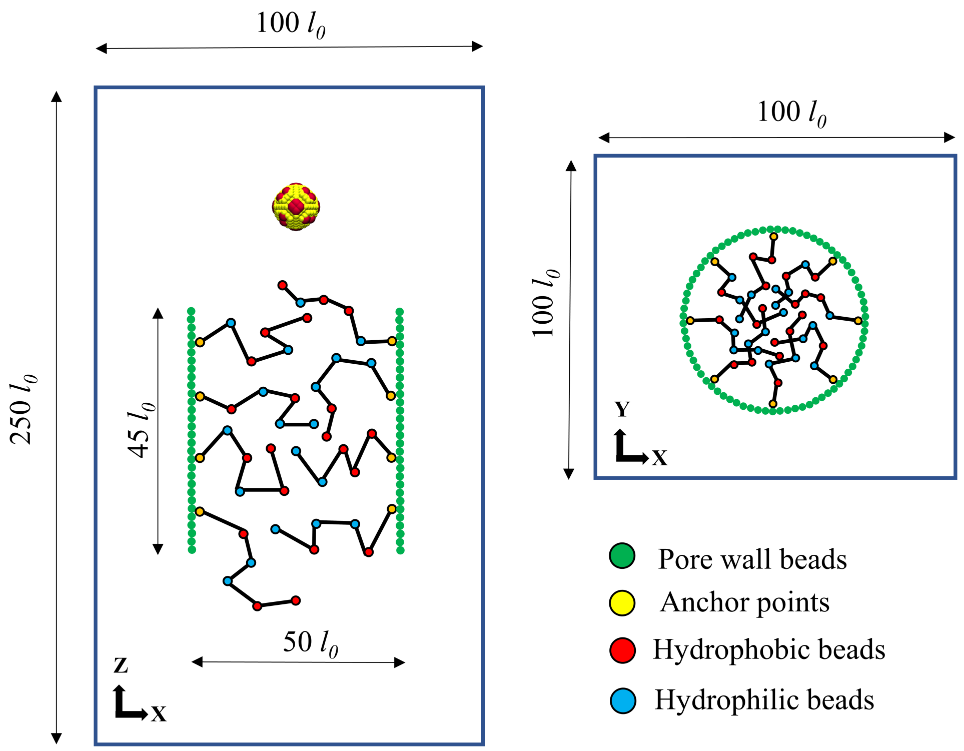


**Figure S1.** Schematic of simulation setup showing the simulation box dimensions and the coarse-grained description of both NPC and tracer.

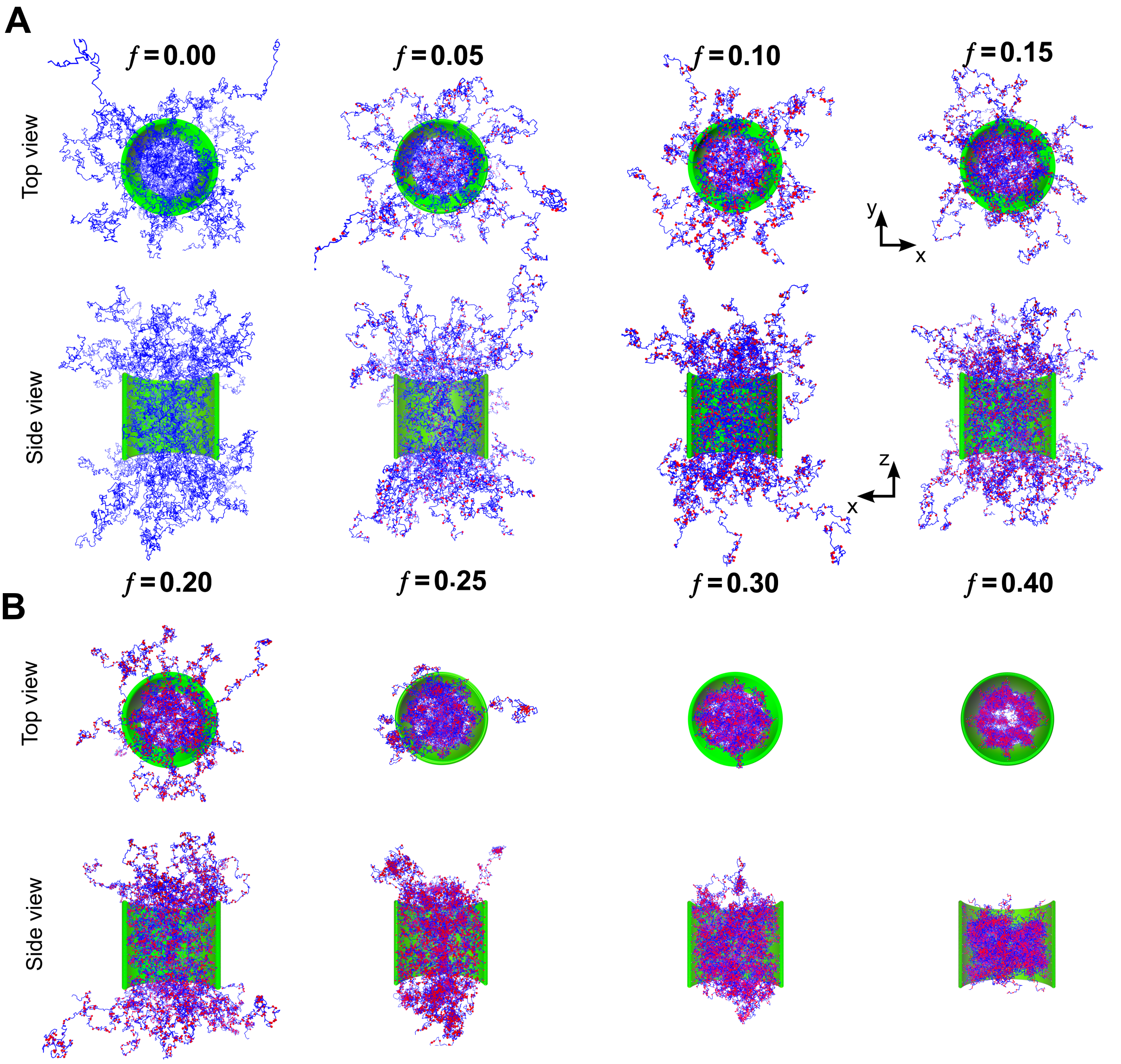


**Figure S2:** Simulation snapshots showing top and side views of equilibrium brush structures for the model NPC corresponding to (A) $f = 0, 0.05, 0.10, 0.15,$ and (B) $f = 0.20, 0.25, 0.30, 0.40$.

#### S3. Stillinger algorithm for identifying clusters

FG-beads (type 2 beads) in the polymer brush formed dynamic crosslinks due to hydrophobic interactions. Collectively, these crosslinks gave rise to the formation of clusters inside the NPC. The Stillinger algorithm was used in the current work to identify and quantify the formation of clusters inside the cylindrical NPC pore (13, 14).

Consider a system of N particles that interact with each other through a short-range pair potential, $V(r)$. An effective interaction distance, $b$, can be defined around each sphere within which pair interaction are significant enough to result in clustering of particles. Overlaps between neighboring spheres can then be obtained by drawing a sphere of radius $b/2$ around every particle. As shown in Figure S3, overlapping spherical regions define a cluster, $C$, constituted of all overlapping particles. If particle $i$ belongs to a cluster, $C$, and $r_{ij} < b$, then $j$ also belongs to C.

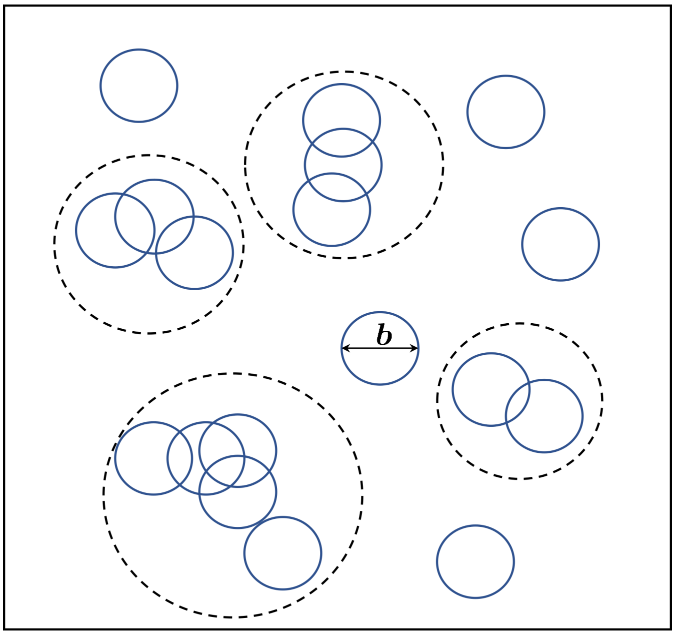


**Figure S3:** Cluster identification algorithm. Overlapped particles of radius b/2 forming clusters. 18 particles shown in configuration shows 1 cluster consisting of 2 particles, 2 clusters consisting of 3 particles and 1 cluster of 5 particles.

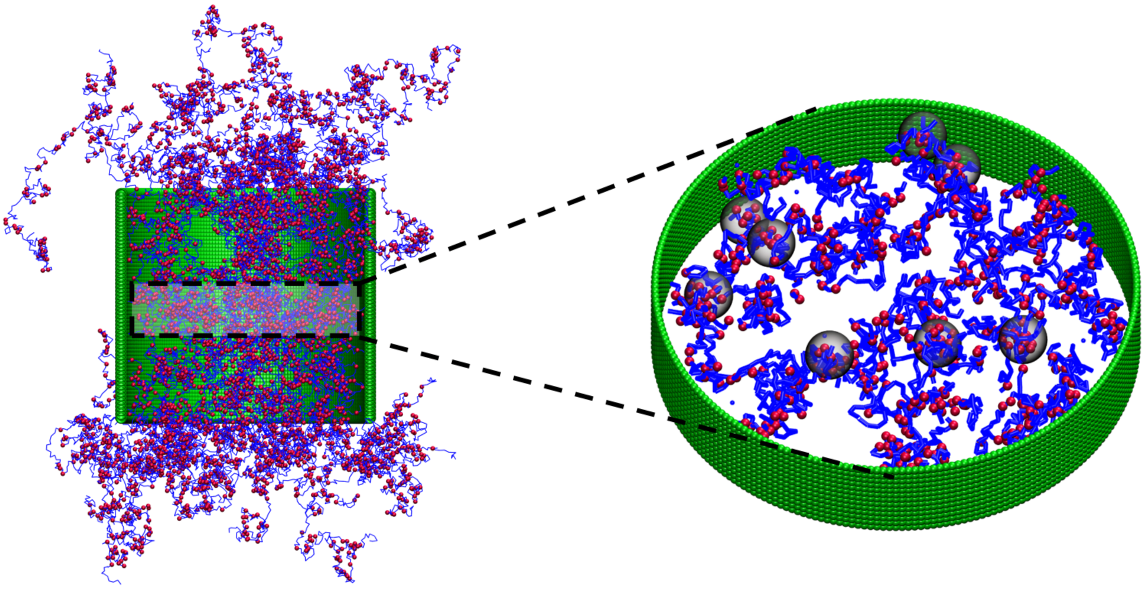

**Figure S4:** Snapshot of cluster formation inside the pore for *f* = 0.2. A part of the brush from only the middle section of the pore (left) is taken to highlight the formation of hydrophobic clusters in the region. Clusters are represented as transparent spheres containing crosslinked hydrophobic beads (right).

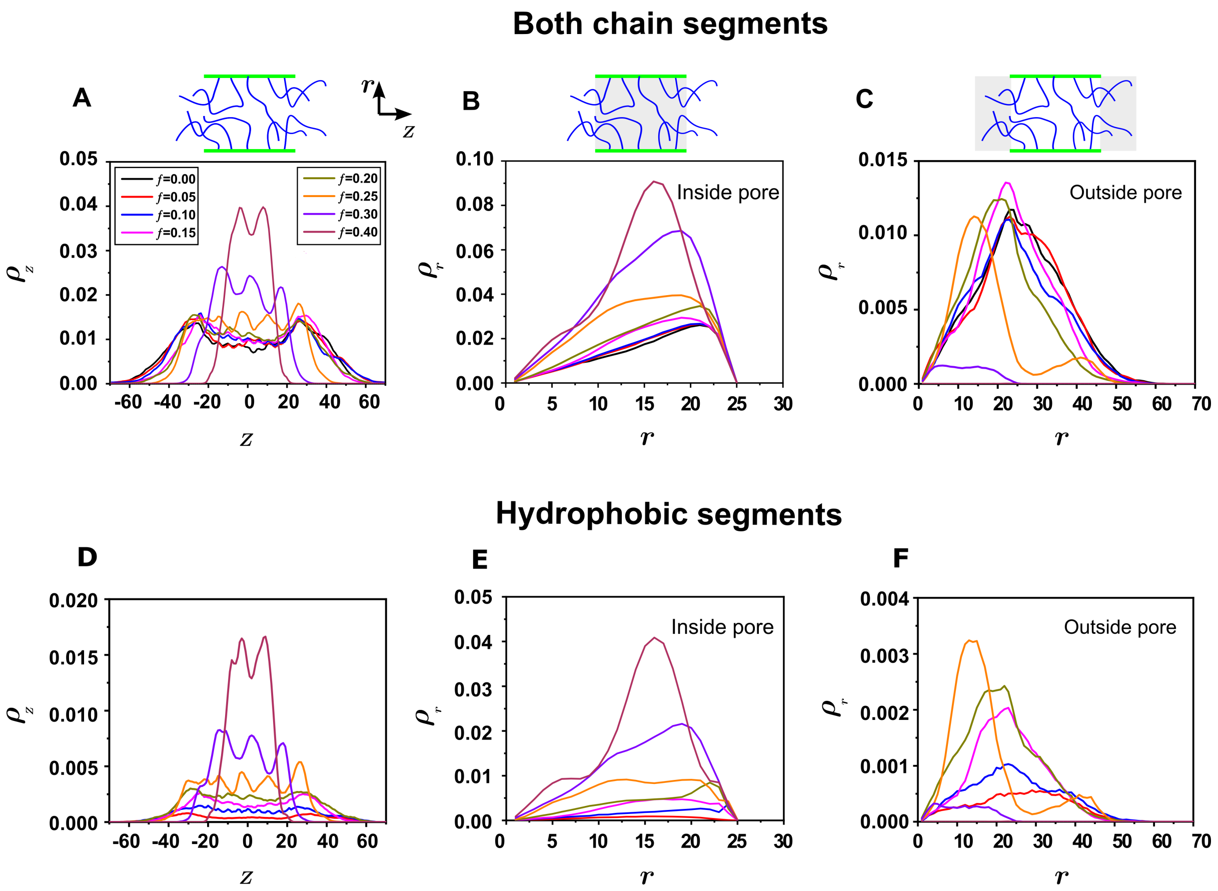


**Figure S5:** Plots showing equilibrated density distributions of brush segments calculated by considering, (A – C) both hydrophobic and hydrophilic segments, and (D – F) only hydrophobic segments. Density distributions were plotted along the (A, D) axial direction of the NPC, (B, E) radial direction of the NPC (inside the cylindrical region), and (C, F) radial direction of the NPC (outside the cylindrical region).

### S4. Tracer trajectories for an inert tracer ($d_t$ = 6 nm)

Nearly all trajectories simulated for inert tracers, $d_t = 6$ nm at $F_t$= 0.8 pN, were rejected by the NPC for all values of $f$ (Figure S5 of the Supporting Information). The scale of thermal force in the simulation **is**, $k_B T/l_0 \approx 4$ pN. Hence, $F_t$= 0.8 pN was much weaker compared to the Brownian forces on tracers which prevented tracers from entering the NPC. In addition, the brush also offered a large entropic barrier to tracer entry at low $f$ because of its significant expanse outside the pore. In contrast, all tracer trajectories of 6 nm inert particle translocated successfully at $F_t$= 4 pN for every hydrophobic fraction (Figure S6 of the Supporting Information). At low *f*, the tracer was small enough to pass through the open structure of the NPC. For $f \geq 0.3$, the tracer size was small enough to pass through the gaps in the tightly bound network structure. Typical translocation times of ≈ 0.5 ms were observed for $F_t$= 4 pN.

At an intermediate force of $F_t$= 2 pN, some differentiation in tracer response to the FG hydrophobic fraction was observed (Figure S7 of the Supporting Information). A simple back-of-the envelope calculation based on experimental measurements of concentration gradients shows that a value of $F_t$= 2 pN is appropriate in the biophysical (3). Most of the 12nm tracer trajectories for $f \geq 0.25$ were either rejected or trapped by the NPC indicating a signature of pore selectivity with respect to the FG hydrophobic fraction, *f.* However, the motion of a 6 nm tracer corresponded to the case of passive translocation through the NPC not requiring transport receptors. In contrast, the larger 12 nm tracer corresponded to the case of facilitated transport across the NE aided by transport receptors (NTR/Kaps). This is supported by reports in the literature where (3–5).

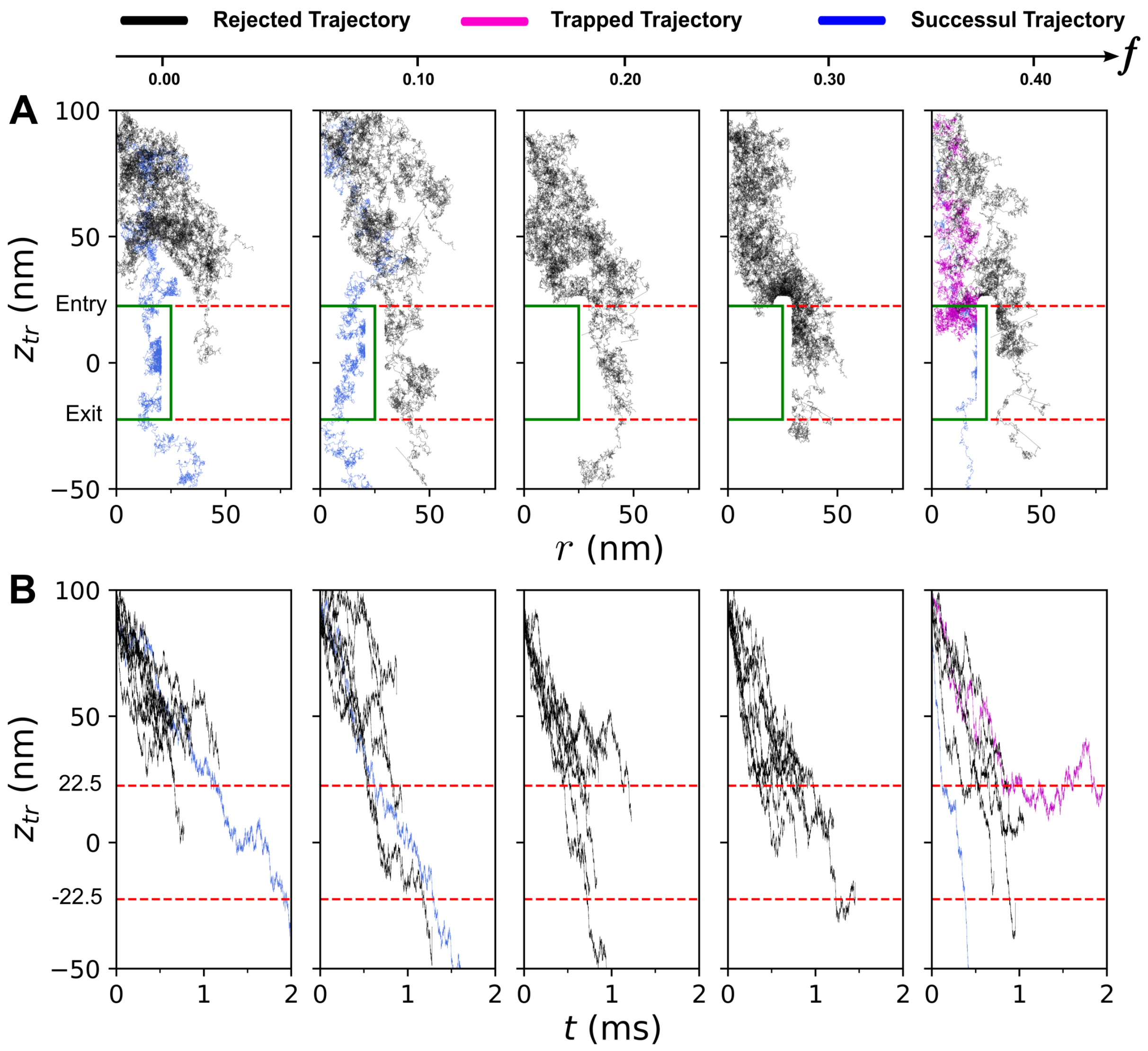


**Figure S6:** Inert tracer trajectories from ten independent simulations shown for different values of $f$ corresponding to $d_t$= 6 nm, $F_t$ = 0.8 pN. The simulated trajectories were classified into three types, *successful* (blue), *trapped* (pink) and *rejected* (black). Successful trajectories corresponded to tracers "successfully" entering the NPC and exiting from the other side. (A) Tracer paths during the simulations represented as $z_{tr}$ *vs* $r$ plots, where $z_{tr}$ is the $z$-coordinate of the tracer, and $r$ is tracer radial coordinate. (B) Plots showing the variation of $z_{tr}$ with time, $t$.

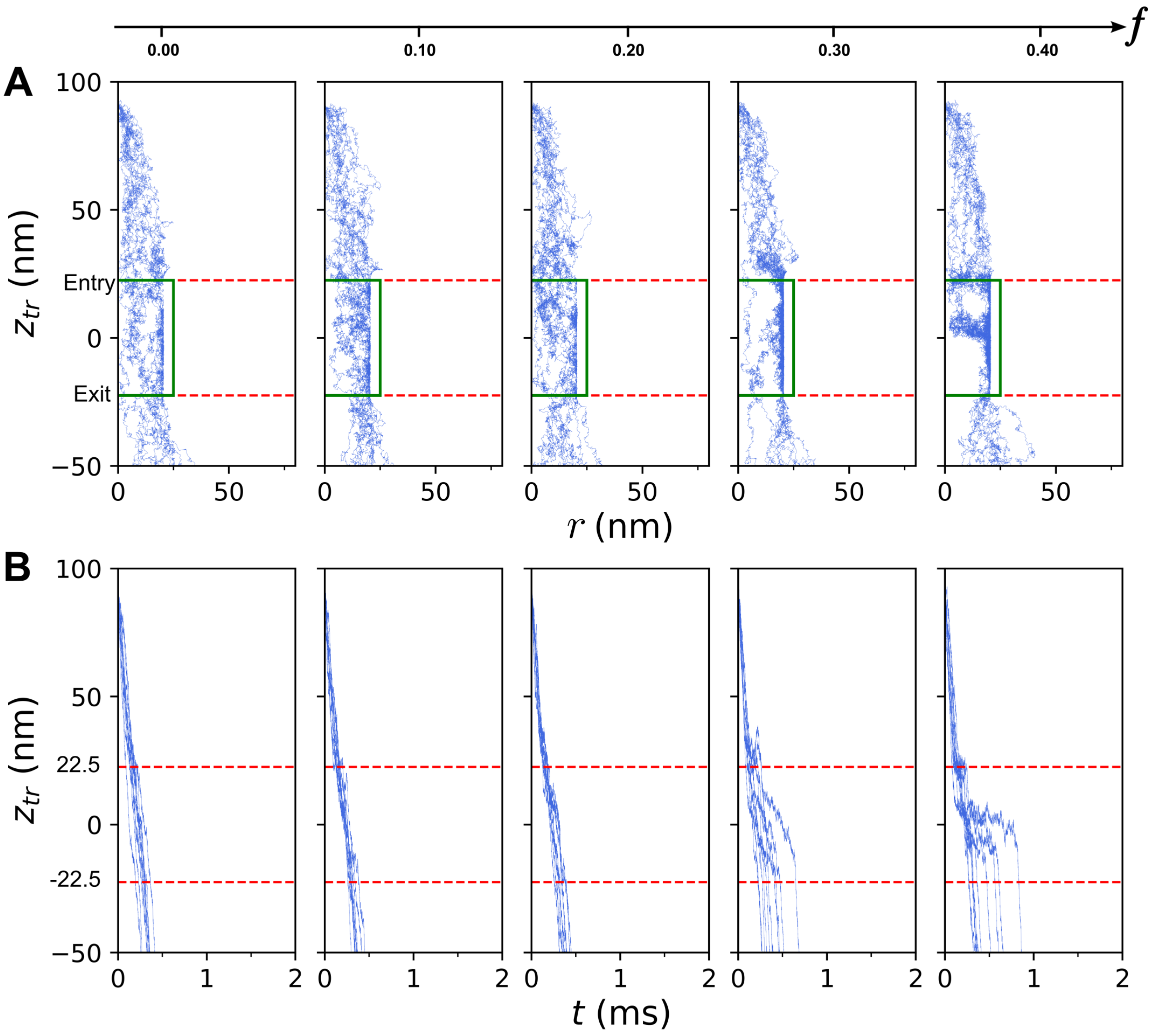


**Figure S7:** Inert tracer trajectories from ten independent simulations shown for different values of $f$ corresponding to $d_t$= 6 nm, $F_t$ = 4.0 pN. (A) Tracer paths during the simulations represented as $z_{tr}$ *vs* $r$ plots, where $z_{tr}$ is the $z$-coordinate of the tracer, and $r$ is tracer radial coordinate. (B) Plots showing the variation of $z_{tr}$ with time, $t$.

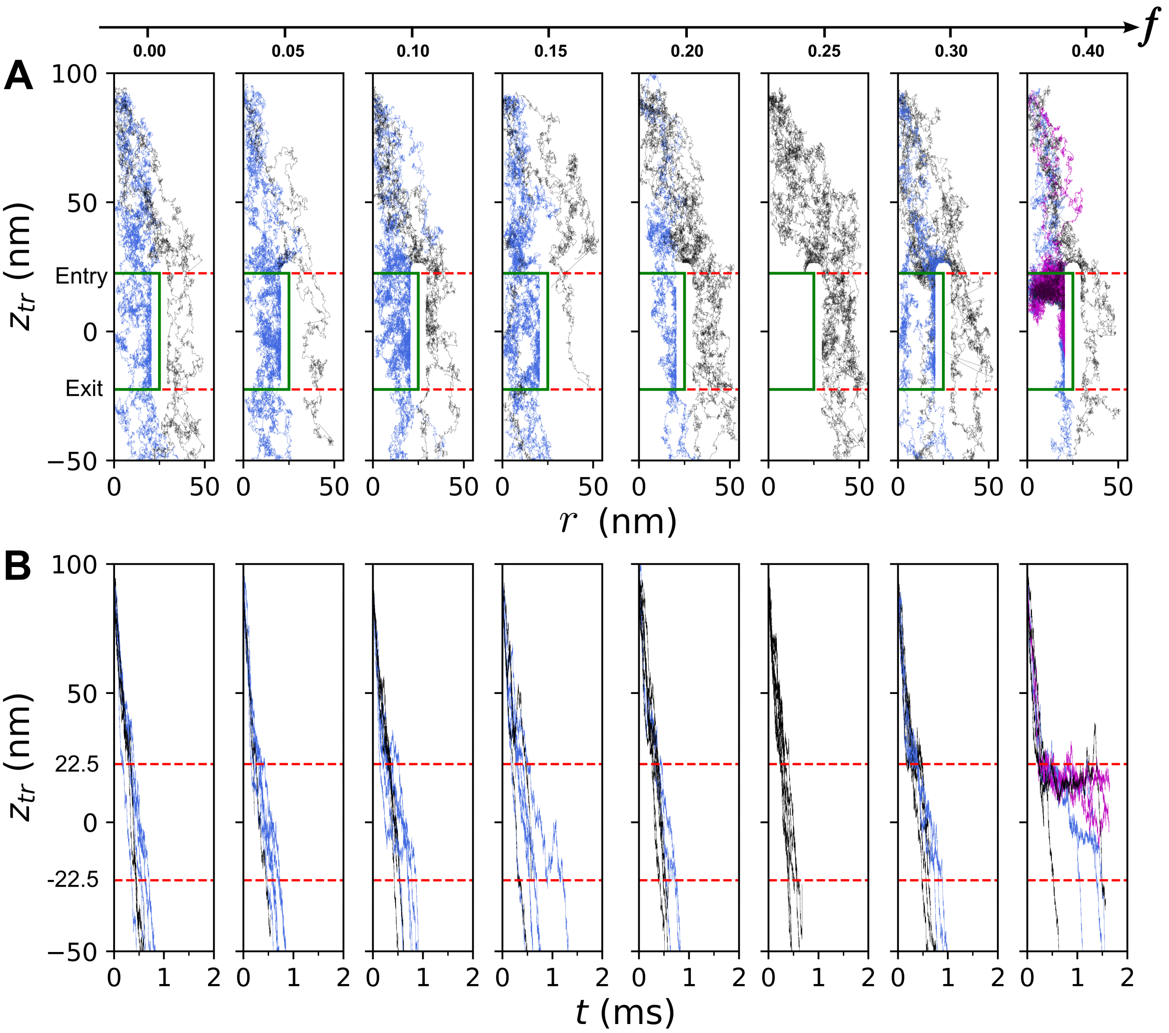


**Figure S8:** Inert tracer trajectories from ten independent simulations shown for different values of $f$ corresponding to $d_t$= 6 nm, $F_t$ = 2.0 pN. (A) Tracer paths during the simulations represented as $z_{tr}$ *vs* $r$ plots, where $z_{tr}$ is the $z$-coordinate of the tracer, and $r$ is tracer radial coordinate. (B) Plots showing the variation of $z_{tr}$ with time, $t$.

### S5. Tracer trajectories for inert and patchy tracers ($d_t$ = 12 nm)

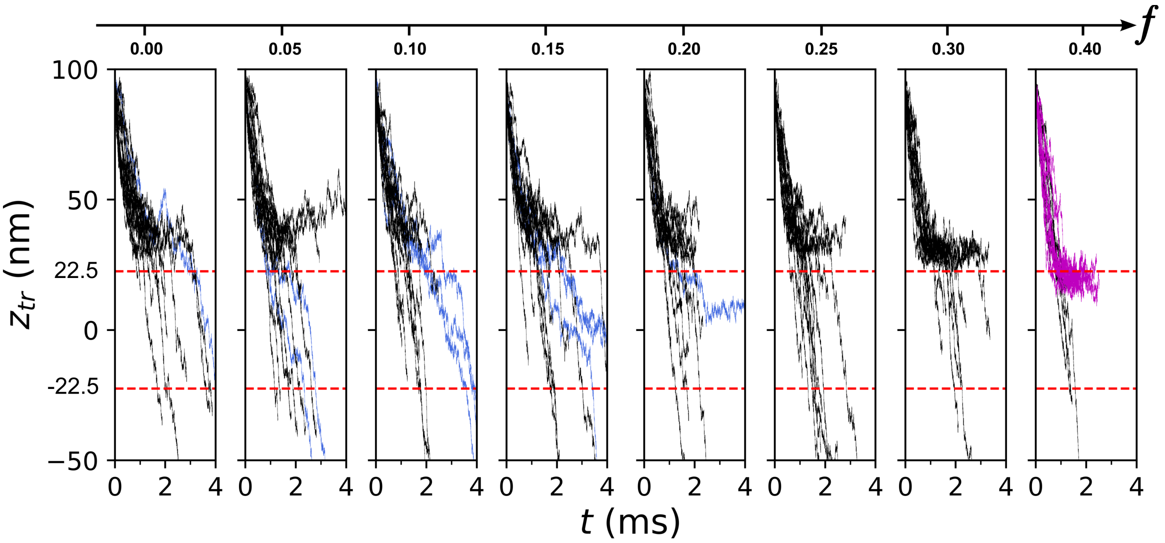


**Figure S9:** Inert tracer trajectories from twenty independent simulations shown for different values of $f$ corresponding to $d_t$= 12 nm, $F_t$ = 2.0 pN. Plots show the variation of $z$-coordinate of the tracer, $z_{tr}$, with time, $t$.

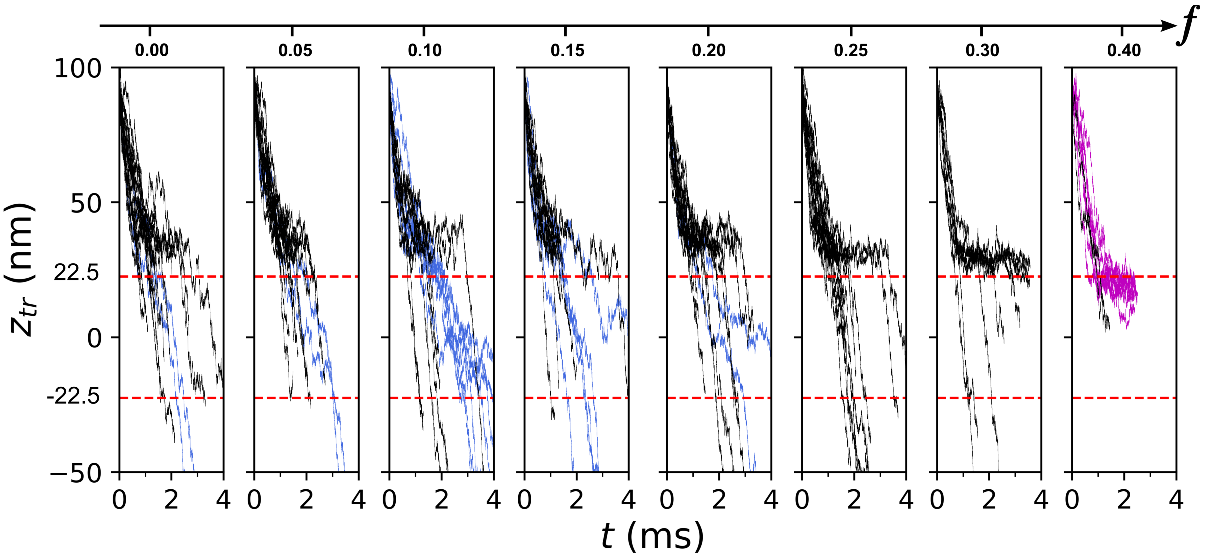


**Figure S10:** Patchy tracer trajectories from twenty independent simulations shown for different values of $f$ corresponding to $d_t$= 12 nm, $\epsilon_{26}$ = 1.5, $F_t$ = 2.0 pN. Plots show the variation of $z$-coordinate of the tracer, $z_{tr}$, with time, $t$.

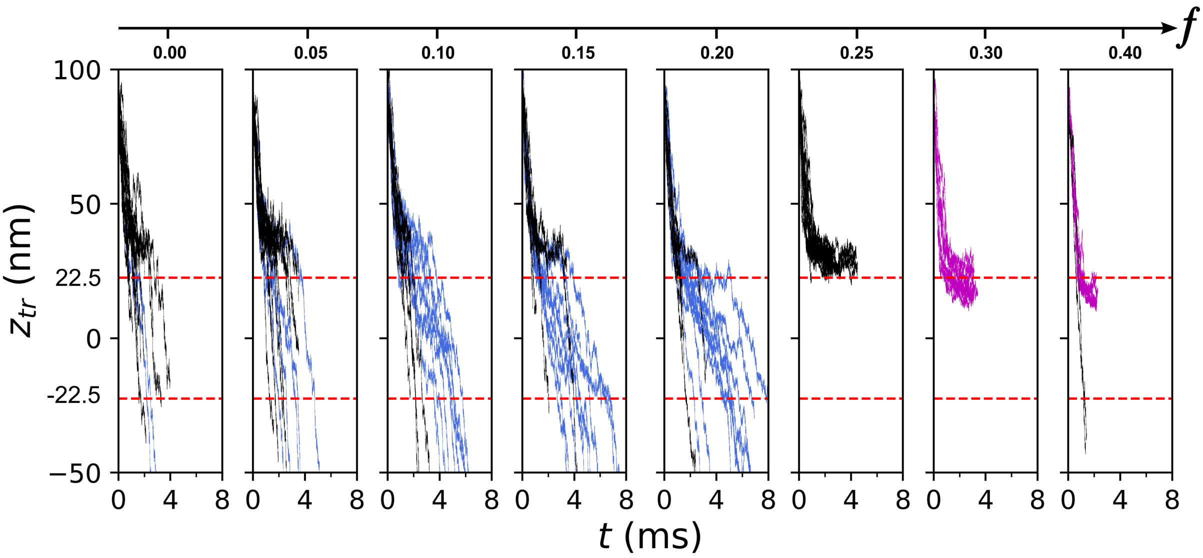


**Figure S11:** Patchy tracer trajectories from twenty independent simulations shown for different values of $f$ corresponding to $d_t$= 12 nm, $\epsilon_{26} = 2.0$, $F_t = 2.0$ pN. Plots show the variation of $z$-coordinate of the tracer, $z_{tr}$, with time, $t$.
